# Central versus peripheral neural control of a coordinated walking pattern in *Drosophila*

**DOI:** 10.64898/2026.04.29.721658

**Authors:** Neha Sapkal, Divya S. Kumar, Suhas Sunke, Nino Mancini, Jason Pitchford, Kazuma Murakami, Salil S. Bidaye

**Affiliations:** Max Planck Florida Institute for Neuroscience.

## Abstract

Walking involves coordinate rhythmic movements at every joint of every leg. Central pattern generator (CPG) circuits in the spinal or ventral nerve cord provide such a rhythmic drive to all the legs^1–3^. In turn, each leg provides rhythmic sensory feedback through peripheral proprioceptive neurons^2,4–6^. Disentangling contributions from these two rhythmic drives, has been a long-standing hurdle in uncovering both the structure and function of neural-circuits governing the generation of a coordinated walking output^2,3,7^. Here, using the highly tractable *Drosophila* model and a novel sensory-deprivation paradigm, we uncovered central and peripheral neural pathways underlying walking pattern generation. We provide evidence that each leg is governed by its own CPG module with an inherent cycle period that is unmasked when proprioceptive feedback is reduced. We find that contact driven load inputs and descending brain inputs are critical for coordinating the intra-leg movements and shaping the microstructure of a single leg’s step-cycle. We show that central coupling pathways underlie inter-leg coordination and that proprioceptive inputs and descending brain commands can flexibly modulate this coupling in a task-specific manner. We identify co-stimulation of specific descending neurons as a mechanism for speeding-up this walking rhythm. Finally, by constraining connectome search^8^ based on the empirical results, we identify putative neural-circuit motifs underlying generation of a coordinated six-legged walking pattern.

## Main

More than a century ago, T. G. Brown proposed that a basic walking pattern could be generated purely by central pattern generating (CPG) elements in the spinal cord^1,9^. Brown’s CPG concept received push-back from the predominant view of the time that rhythmic proprioceptive feedback is the primary driver of stepping motor patterns through reflexive actions^10–12^. It was only much later that the CPG concept was revived, initially using invertebrate model systems by researchers studying other motor rhythms^13–15^. The first evidence of a central neuron instrumental to walking rhythm generation was provided through intracellular recordings in the cockroach nerve cord^16^. This was followed by similar discoveries in other insects^17–19^. Alongside these discoveries in invertebrates, mammalian locomotion research across several species, supported the idea that each leg is controlled by its own spinal CPG module^7,20,21^. Based on these empirical data, currently, there exist a few mutually non-exclusive theoretical models of how a walking CPG network drives a coordinated motor output^22–25^, however the biological correlates of these models are still being uncovered. A recent, connectome-constrained modeling study in *Drosophila* provides a putative single-leg stepping CPG circuit, where we know each neuronal type and all the connectivity^26^, yet it remains to be functionally verified.

In parallel with progress in uncovering the walking CPGs, a large body of work using both insect^2,5,27–30^ and mammalian models^4,6,31–34^ showed that sensory feedback from the stepping legs is indeed instrumental in setting up a proper walking pattern which includes both intra-leg motor coordination, as well as inter-leg coordination. In insects it has been proposed that this sensory-feedback contribution is walking-speed dependent^5,28,35–38^. Stick insects tend to walk slowly on unpredictable terrains and show strong influence of sensory feedback compared to insects like cockroaches that typically run at high-speeds on flat terrains^5,28,38,39^. However, how specific sensory feedback pathways interact with central CPG circuits is still unclear.

One reason for this knowledge gap, especially in case of insects, is the lack of reproducibility and specificity in inducing similar walking rhythms in intact and deafferented animals. Previous efforts to isolate central versus peripheral contributions to pattern generation in insects have typically relied on comparing recordings from “real walking” intact or semi-intact insects to recordings from deafferent preparations and isolated nerve cords that are induced into a “fictive walking” state by application of muscarinic agonist, pilocarpine^17,18,38^. While these studies have been highly impactful^2^, pilocarpine application (non-specific activation across several neuronal types) produces variable outcomes across preparations, drug concentration and insect species. More importantly, it is unclear how similar the drug-induced “fictive walking” is to “real walking”^17–19^. In addition, this approach does not provide obvious targets of where the CPGs might be and relies on “blind” electrophysiological recordings in fictive walking preparations^2^.

The mammalian walking CPG studies also initially relied on pharmaco-activation of deafferented spinal cords^20^. However, in later years, the work benefitted from the ability to reliably induce fictive walking patterns in deafferented preparations by electrically or optogenetically stimulating MLR (mid-brain locomotor region) or spinal-cord^20,21,40–43^. This has led to enormous progress in the field, to the point where some genetically defined CPG candidate-neurons are now characterized in the mouse spinal cord^44^. Yet, robust multi-limbed walking patterns remain difficult to reproducibly stimulate in deafferented preparations^20^. More importantly, connectivity within identified CPG neurons and how sensory feedback pathways connect with these CPG motifs, is still unresolved^20^. These aspects are critical for mechanistic understanding of how a coordinated multi-limbed walking pattern is generated.

Inspired by these studies, we previously identified MLR analogous brain neurons that control walking initiation in the fruit fly *Drosophila melanogaster*^45^. In a subsequent study, by pinpointing where GABAergic halting neurons converge on outputs of these walking initiation neurons, we identified descending neurons (DNs, that project from the brain to the nerve cord) critical for walking initiation^46^. In this work, by designing a novel sensory deprivation paradigm coupled with optogenetic stimulation of these critical walking promotion DNs, we could induce different walking patterns with extreme precision and reproducibility across both intact and deafferented, decapitated preparations. This approach allowed overcoming persistent challenges faced by the previous studies and resulted in an unprecedented view into central versus peripheral contributions to walking pattern generation. Guided by these highly quantitative empirical data, we parsed the available *Drosophila* ventral nerve-cord connectomes^8,47,48^ to reveal the underlying neural circuitry for both intra-leg as well as inter-leg coordination.

### Evidence for a distributed CPG based control of stepping rhythms

Previously, we identified three critical walking promotion DNs in *Drosophila*, that on activation drive walking: MDN drives backward walking^49^, whereas DNg100 (also referred to as BDN2^46^) and DNg97 (also referred to as oDN1^46^) drive forward walking^46^. Remarkably, optogenetic stimulation (with UAS-CsChrimson^50^) of the severed axons of these DNs in decapitated flies, induces backward or forward walking, respectively (Fig. 1a-b, Extended Data Fig. 1a-c). This shows that the nerve cord outputs of these neurons are sufficient to drive walking and therefore provide a perfect platform for addressing how cord circuits generate a coordinated walking pattern. The use of decapitated preparations with optogenetic stimulation of DNs allows us to study walking output without any sensory or internal-state influence coming from the brain. We have further devised a set of paradigms that progressively reduce sensory feedback from the stepping legs to decipher how much each proprioceptive element contributes to the optogenetically triggered walking pattern. This allows us to systematically analyze each genotype across four different conditions (Fig. 1c):

**Fig. 1:**
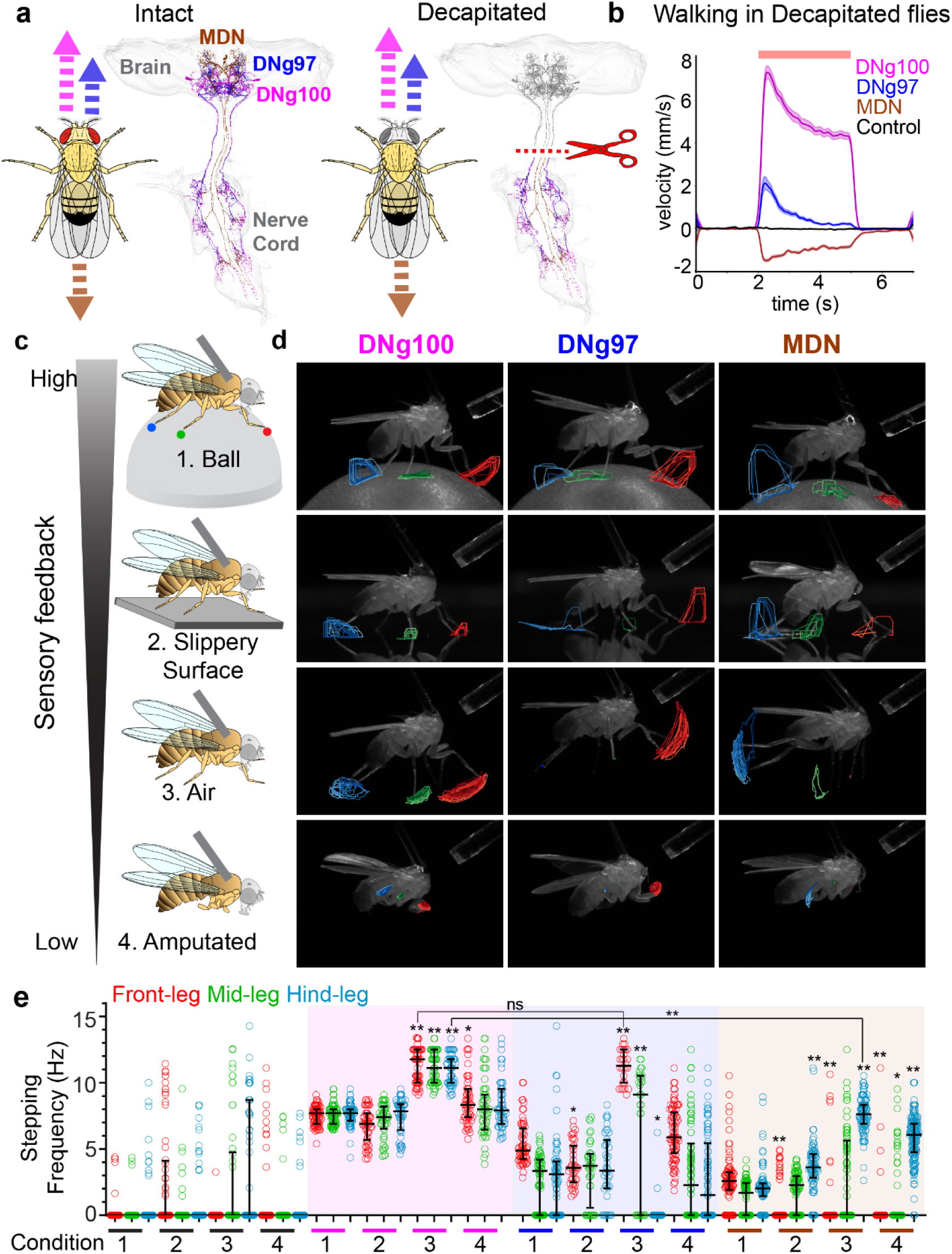
Empirical evidence for distributed CPG driven walking: **a.** Anatomy of critical walking promotion descending neurons (DNs) and schematic of decapitated condition. **b.** Forward walking velocity (mean ± s.e.m) of tethered decapitated flies during optogenetic stimulation (red bar) of respective DNs (colored) and controls (black). **c.** Schematic of different conditions for progressively reducing proprioceptive contribution to walking in decapitated flies. **d.** Example video frames with overlaid tarsal tip tracking data of each DN stimulated flies stepping in conditions from c. **e**. Stepping frequency (median ± interquartile, each point is a trial) for each DN stimulated and control flies in each condition. n = 60 to 70 trials across 6 to 7 flies per genotype and condition. Kruskal-Wallis test followed by Dunn’s multiple comparison with respective leg’s condition-1 data within same genotype. For condition-3 (air) dataset we also performed post-hoc comparison across front leg data of DNg100 and DNg97 and across hind-leg data of DNg100 and MDN (**p <0.001, * p< 0.05, ns: non-significant is only written for one highlighted case). (also see Supplementary Videos 1-3). Supplementary Table 1 shows full experimental genotypes and exact sample sizes.

#### Condition-1

##### Ball

This is the conventional preparation for tethered (decapitated) walking *Drosophila* on an air-supported ball (spherical treadmill)^51–53^. The legs step on the ball and are mechanically coupled through the rigid substrate and body and all proprioception is intact.

#### Condition-2

##### Slippery surface

The legs are mechanically decoupled (movement of one leg does not exert mechanical forces on the other leg) because the tethered fly steps on a slippery surface (hydrophobic coverslip). However, most proprioceptive inputs are still intact.

#### Condition-3

##### **Air**-stepping

Tethered fly is suspended in air with no substrate to grab on to. As a result, there is no mechanical coupling and no ground-contact induced load inputs. Leg movements will still recruit some proprioceptors like the joint-movement sensors (Chordotonal Organs^4^), however load sensors (“Campaniform Sensilla, CS”^4^) are non-functional.

#### Condition-4

##### Air-stepping with amputated legs

Same as condition-3 but now all legs are amputated at the mid-point of the femur such that the major movement sensor (Femoral Chordotonal Organ, FeCO^4^) is no longer present.

As we go from Condition-1 (“Ball”) to Condition-4 (“Amputated”), the peripheral contribution (sensory feedback from legs) gradually decreases to a point where in Condition-4, the preparation is reduced to cord-circuits controlling stumps of legs with very little (if any) sensory feedback. We used previously established highly specific split-Gal4 reagents^46,49^ to optogenetically stimulated the severed axons of DNg100, DNg97 and MDN in decapitated flies, across all conditions and recorded high-speed videos (200 Hz) from eight camera angles, as described in our previous work^46^. We then optimized a previously established 3D-leg kinematics analysis pipeline^46^ to track and analyze every joint of every leg of each experimental and control fly. This also required developing a new proofreading tool to ensure proper 3D tracking across conditions (released to the community as a part of this work, see Methods). We found that while control decapitated flies do not show stepping in any of the conditions, the DN stimulated decapitated flies, showed rhythmic stepping across all four conditions (Fig. 1d, Extended Data Fig. 1c and Supplementary Videos 1-3). This indicates that the central neural circuits in the nerve cord, triggered by the DN, are sufficient to generate stepping rhythms.

In case of DNg100 stimulation, all six legs show rhythmic forward-stepping across all four conditions (Fig. 1d-e, Extended Data Fig. 1c, Supplementary Video 1). Stepping frequency was higher during “Air-stepping” compared to other conditions, likely due to lack of ground-contact. However, air-stepping was even faster than stepping in amputated leg-stumps, which could be potentially attributed to lack of FeCO inputs or some tissue damage caused by amputation. To distinguish between these two possibilities, we repeated the above experiment while optogenetically silencing FeCO (co-expressing anion channelrhodopsin GtACR1^54,55^ using an iav-LexA reagent^56^, Extended Data Fig. 1d). Silencing FeCO caused the stepping frequency to reduce even during non-amputated air-stepping bringing it down to amputated levels (Extended Data Fig. 1e). This confirms that the “amputated” condition is a highly sensory-deprived state that removes FeCO-derived sensory input in addition to load inputs and confirms that FeCO inputs are necessary for increased stepping frequency. The only remaining functional proprioceptors in this condition are proximal joint limit-detectors (hair-plates)^57^. To address if proprioceptive feedback from these proximal sensors is required for observed stepping, we immobilized proximal joints of front-legs by gluing these joints and quantified movement at the distal Femur-Tibia (Fe-Ti) joint while stimulating DNg100 in decapitated flies. Even in this highly unnatural condition (legs are immobilized in a raised state that never occurs during regular walking), we observed robust Fe-Ti oscillations indicating that sensory feedback from these proximal hair-plates is not a requirement for DNg100 induced stepping (Extended Data Fig 1f-h and Supplementary Video 4). Similar to amputated condition, the stepping frequency was lower than intact leg “air-stepping” suggesting a potential role for hair-plates in controlling stepping frequency.

In contrast to DNg100, stimulation of DNg97 and MDN produced pronounced differences across the four sensory deprivation conditions and across legs. DNg97 stimulation drove stepping rhythms across all six legs in the ball-condition. However, as mechanical coupling and sensory feedback was reduced, stepping was predominantly seen only in front legs, and to a smaller extent in mid-legs (Fig. 1e, Extended Data Fig 1c and Supplementary Video 2). This shows that DNg97 actively recruits only front-legs. Also, similar to DNg100, the stepping frequency for air-stepping in front-legs was much higher compared to other conditions and reached the same level as DNg100 air-stepping (Fig. 1d-e).

MDN stimulation showed a reversed phenotype compared to DNg97 stimulation. MDN drove backward stepping across all six legs in ball-condition (Fig. 1d-e). However, when sensory feedback was reduced MDN predominantly drove rhythms in hind legs (Fig. 1d-e, Supplementary Video 3), in line with previous studies that showed hind legs are the primary targets of MDN stimulation^49,58,59^. Flies, like most other animals, typically walk backwards much slower than forwards^49^. Consistent with this, the median stepping frequency of MDN stimulated flies walking on the ball was very low (2 Hz for hind legs). Remarkably, the median air-stepping frequency of MDN stimulated flies’ hind legs was much higher (7.6 Hz) and approached the forward air-stepping frequencies (∼11Hz) observed for DNg100 and DNg97.

Taken together, we found that in sensory-deprived conditions different DNs drive stepping across different sets of legs (DNg100: all six legs, DNg97: front legs, MDN: hind legs) at similar stepping frequencies (Fig. 1e). These results suggest the existence of a distributed set of common CPG modules (at least one CPG module per leg) that are recruited by the DNs in a segment specific manner. The oscillation of distal joints while proximal joints are glued and oscillations of proximal joints while distal joints are amputated, shows that each leg-joint can be directly driven into rhythmic movements by CPGs without feedback from other joints.

### Central circuits in the nerve-cord drive a symmetric step-cycle that is structured by proprioceptive feedback and descending inputs

To address intra-leg coordination, we analyzed the micro-structure of the optogenetically induced step-cycles of each leg in the decapitated flies. Stepping was highly reproducible for the “actively” stepping legs in each case described in Fig. 1c-d (DNg100: all legs, DNg97: front-legs and MDN: hind-legs). We could therefore analyze the averaged step cycle (Fig. 2a-e, Extended Data Fig. 2a-b, see Methods) across each condition and stepping direction (forward versus backward) to obtain a comparative view. For brevity, in Fig. 2a-e we focus on the movement of a forward stepping front leg (DNg100 stimulated), a forward stepping hind leg (DNg100 stimulated) and a backward stepping hind leg (MDN stimulated), and we report all other leg movements in Extended Fig. 2. By comparing the average trace of each of the main joint angles we can appreciate the similarities and differences across these cases (we ignore tibia-tarsal joint movements for this study because they are small, variable and prone to tracking errors, compared to other joints, see Methods).

**Fig. 2:**
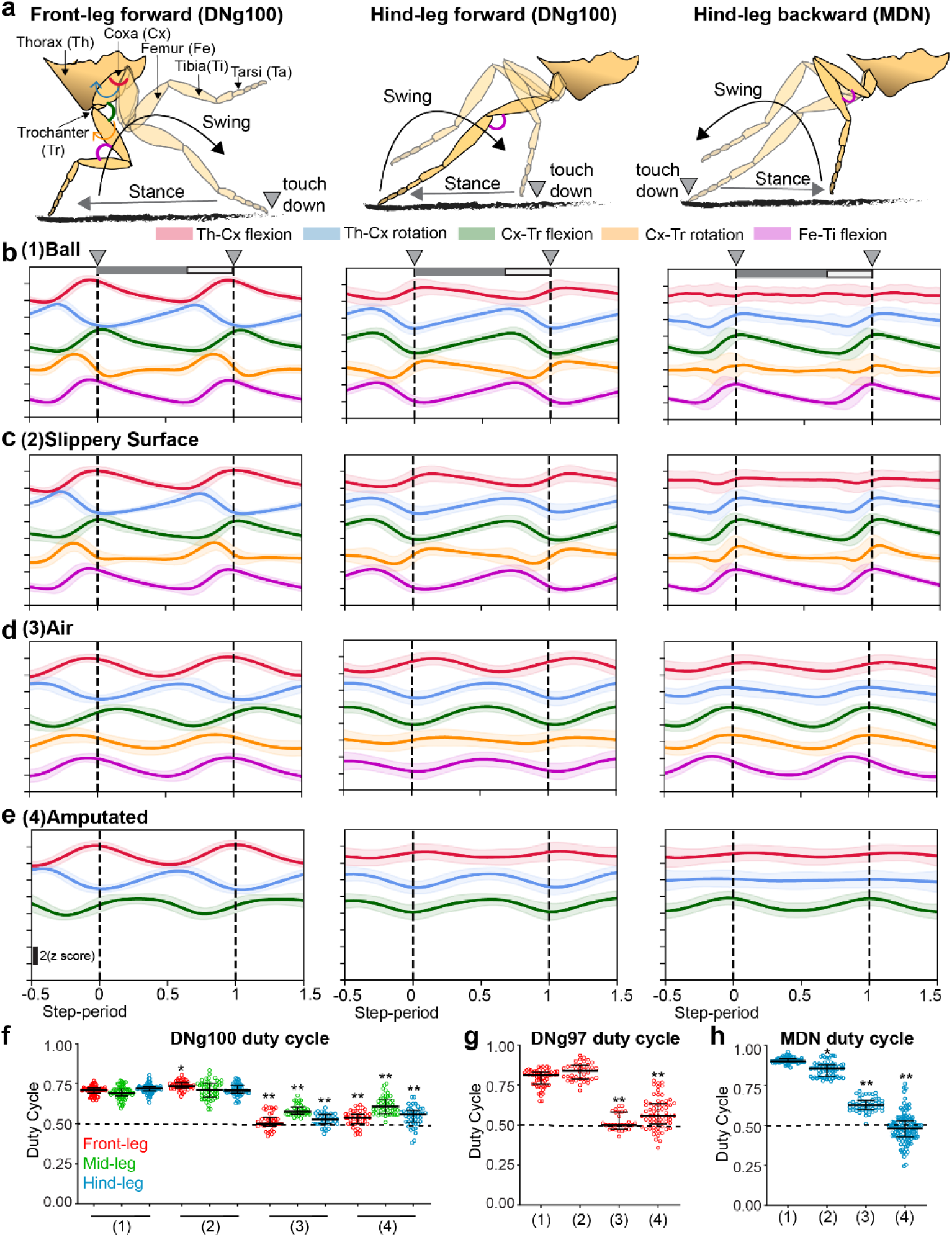
Centrally generated symmetric step-cycle is structured by DNs and sensory feedback: **a.** Cartoon fly leg stepping shown for a forward stepping front leg (left) and hind leg (middle), and a backward stepping hind leg (right). Leg segments are labeled and respective joint angles are depicted and color coded in the left panel. **b-e** Averaged step cycle plots showing each joint angle trace (mean ± s.e.m., color coded as in panel a) for a DNg100 stimulated forward stepping front leg (left) and hind leg (middle) and an MDN stimulated hind leg (right), in case of condition-1 (**b**), condition-2 (**c**) condition-3 (**d**) and condition-4 (**e**). **f-h.** Duty-Cycle (Stance period/step period) for each stepping leg for DNg100 (**f**), DNg97 (**g**) and MDN (**h**) stimulated flies stepping in each sensory deprivation condition from Fig. 1c. n = 28 to 120 trials across 3 to 12 flies. Kruskal-Wallis test followed by Dunn’s multiple comparison with respective leg’s condition-1 data within same genotype. (**p <0.001, * p< 0.05) (also see Supplementary Videos 5). Supplementary Table 1 shows full experimental genotypes and exact sample sizes.

In the ball-condition, across all legs and stepping directions, the step-cycle comprises of a long stance-period (leg on ground) and a short swing-period (leg in air) that leads to a duty cycle (stance-period/total step-period) higher than 0.5 (Fig. 2f-h). Each joint-angle trace has a peculiar shape with obvious asymmetries across swing and stance phase and in case of a DNg100 stimulated fly, these joint-angle traces resemble those of a control head-intact fly spontaneously walking forward (Extended Fig. 2a). Comparing a forward stepping front-leg to a forward-stepping hind leg, it is apparent that while the Thorax-Coxa joint angles retain their profile, the other joint movements (Coxa-Trochanter: Cx-Tr and Femur-Tibia: Fe-Ti) now show a reversed profile. This is expected given how front and hind legs are oriented with respect to the body. Of note, comparing a forward stepping hind leg (DNg100-stimulated) to a backward stepping hind-leg (MDN-stimulated), there is a second reversal in these Cx-Tr and Fe-Ti joint angle profiles so that these joint movements of a backward stepping hind leg now in-turn resemble the movements of a forward stepping front legs. Surprisingly, MDN-driven backward-stepping hind-legs do not have a major contribution from Thorax-Coxa joints. These quantifications suggest that there will be certain similarities and differences in how DNg100 and MDN recruit the premotor circuits that we will revisit in connectome analysis later in the manuscript. In contrast, DNg97 driven front leg movements are kinematically identical to DNg100 driven front leg movements. This kinematic similarity and similarity in stepping frequency in air-condition (Fig. 1e) and amputated-condition, imply that DNg97 and DN100 likely recruit the same premotor circuits for front leg movements. We also noted that DNg97 driven mid and hind-leg movements were highly variable and differ from DNg100 counterparts (Extended Fig. 2a), underscoring the mechanical coupling driven passive recruitment of these movements in the DNg97 ball condition. These data clearly show that DNs directly impact intra-leg coordination to drive changes in stepping direction.

Remarkably, as we move from the ball condition to the slippery-surface condition, even though legs become mechanically decoupled and only “actively” controlled legs now step often in each DN stimulation paradigm, the step-cycles of the actively stepping legs retain the same microstructure as that seen in the “ball” condition (Fig. 2c, Extended Data Fig. 2b). This shows that although mechanical coupling through the substrate impacts which legs step, it does not impact intra-leg coordination.

In contrast, once the walking substrate was removed (air and amputated conditions), the joint movements now show a temporally symmetric pattern (Fig. 2d-e, Extended Data Fig. 2b) with duty-cycle approaching 0.5 across all legs for all DN stimulations (Fig. 2f-h). In these conditions, the joint movement traces lose most of the asymmetries present in the ball-condition and instead now resemble sinusoidal waves (Fig. 2d-e and Extended Data Fig. 2b-c), an output that could naturally result from oscillatory neural activity driven by CPG networks. This shows that presence of substrate (ball or slippery surface) and substrate-driven load feedback (through Campaniform Sensilla, CS^4^), is critical for shaping the micro-structure of a step-cycle and defining swing-stance phase-specific movements.

However, we found that even though there is clear impact on temporal structuring of the joint movements due to sensory feedback, the temporal order in which joints move with respect to the each other is fairly conserved across all conditions. The only manipulation that significantly changed temporal ordering was change in stepping direction brought about by DN stimulation (compare DNg100 stimulated versus MDN stimulated hind leg, Extended Data Fig. 2d).

Thus, the microstructure of a step-cycle is shaped both by descending commands and proprioceptors into a task-specific pattern that defines stepping phase and direction (Supplementary Video 5).

### Descending inputs and proprioceptive feedback modulate central coupling pathways to flexibly tune inter-leg coordination

Insects tend to maintain antiphase relationships between left-right legs of the same thoracic segment (i1-c1, i2-c2, i3-c3, notation: i-ipsilateral, c-contralateral, 1-front, 2-mid, 3-hind, Fig. 3a) as well as between ipsilateral pairs of adjacent thoracic segments (i1-i2 and i2-i3)^60^. At high speeds, most insects tend to exhibit a tripod-like state where three legs (i1-c2-i3) step in-phase with each other and alternate with the complementary tripod^27,35,37,39,61,62^. Our dataset of decapitated walking flies, provided a perfect opportunity to address how central and peripheral inputs contribute to different aspects of inter-leg coordination.

**Fig. 3:**
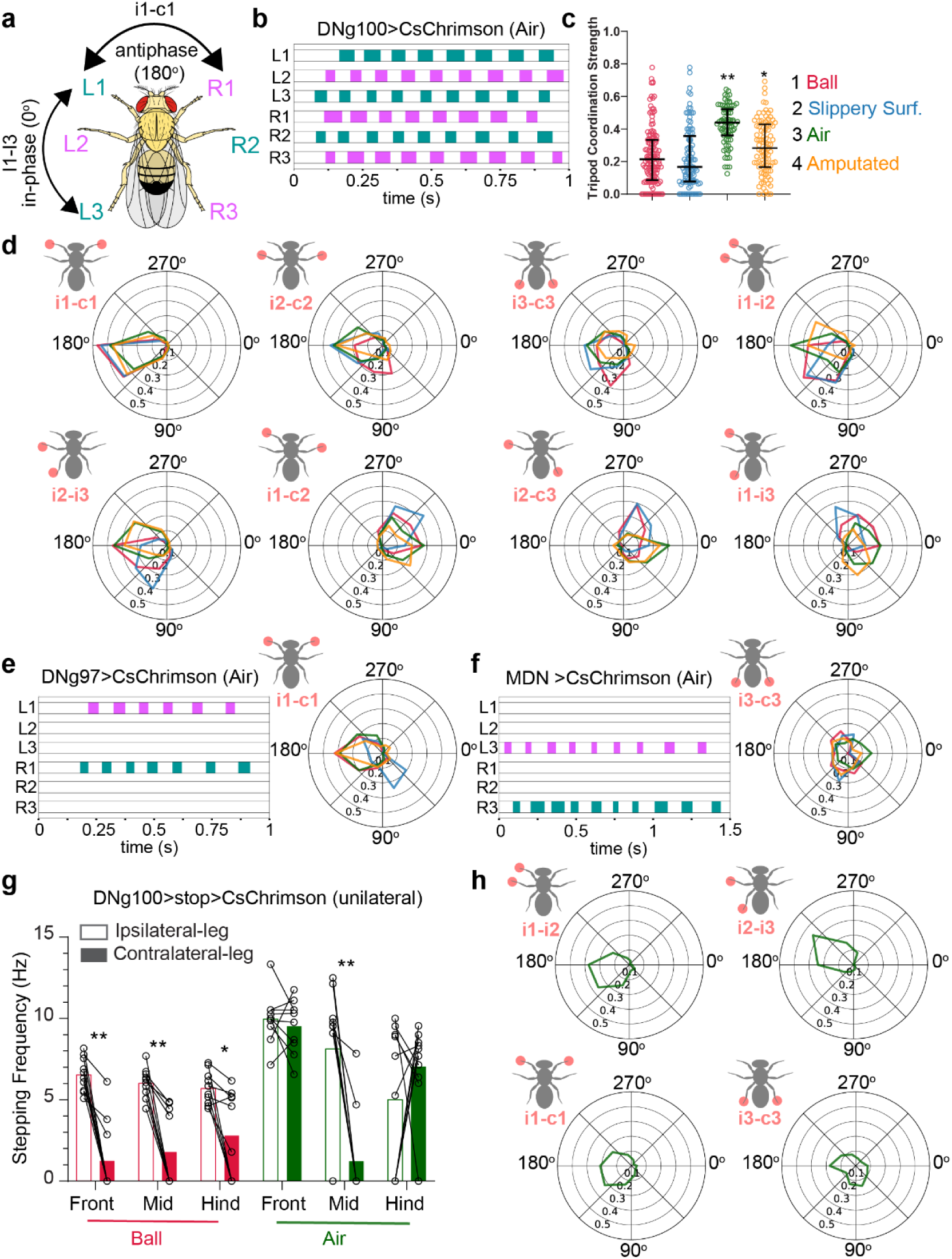
Evidence for central coupling underlying inter-leg coordination: **a.** Schematic of a fly showing inter-leg coordination phases with color coding as in a tripod gait. **b.** Example trial of a DNg100 stimulated fly air-stepping in tripod coordination. **c.** Tripod Coordination Strength (TCS) for DNg100 stimulated flies across all four conditions. n = 38 to 70 trials across 4 to 7 flies. Kruskal-Wallis test followed by Dunn’s multiple comparison with condition-1. **d**. Phase difference plots, between pairs of legs indicated on top left schematic for DNg100 stimulated flies across all four conditions (n= 210-748 phases, see Extended Data Fig. 4 for PLV and statistics). **e-f**. Example stepping pattern (left) and phase difference plot (right) for the pair of actively stepping legs in DNg97 (**e**) and MDN (**f**) stimulated flies across all four conditions (color coding as in **c**). n = 28 to 6 trials across 3 to 8 flies for e, n = 51 to 120 trials across 6 to 12 flies for (**f**). **g.** Stepping frequency of ipsilateral and contralateral legs of unilateral DNg100 stimulated flies across ball and air conditions. Wilcoxon matched-pairs signed rank test for each ipsi- and contra-lateral leg pairs. n = 10 flies, each point is median frequency across trials for each fly (**p <0.001, * p< 0.05). **h.** Phase-difference plot for air-condition from panel g. (also see Supplementary Videos 6-7). Supplementary Table 1 shows full experimental genotypes and exact sample sizes.

We first quantified the “tripodness” of DNg100 stimulated decapitated flies using a stringent parameter, Tripod Coordination Strength (TCS)^37^, that measures degree of overlap across swing phases of ipsilateral front, contralateral mid and ipsilateral hind legs (i1, c2, i3, Fig. 3a-c). We found that DNg100 stimulated flies showed tripod like walking across all conditions with highest TCS in the air condition, implying the existence of central coupling pathways (Fig. 3c, Supplementary Video 6). Both amputated condition (Fig. 3c) as well as FeCO silenced air-stepping (Extended Data Fig. 3a) show reduced TCS implying role of FeCO in inter-leg coordination.

To probe specific inter-leg coordination strengths, we quantified the coupling between each pair of stepping legs across all DN stimulations and conditions by calculating the phase difference between step-cycles of the leg-pairs (Fig. 3d). We also report coupling strength (Extended Data Figs. 3b and 4a-i) as “Phase Locking Value” (PLV^61,63^ that ranges from 0-1 with 0 being no coupling and 1 being perfect coupling, see Methods) corresponding to each phase difference quantification.

**Fig. 4:**
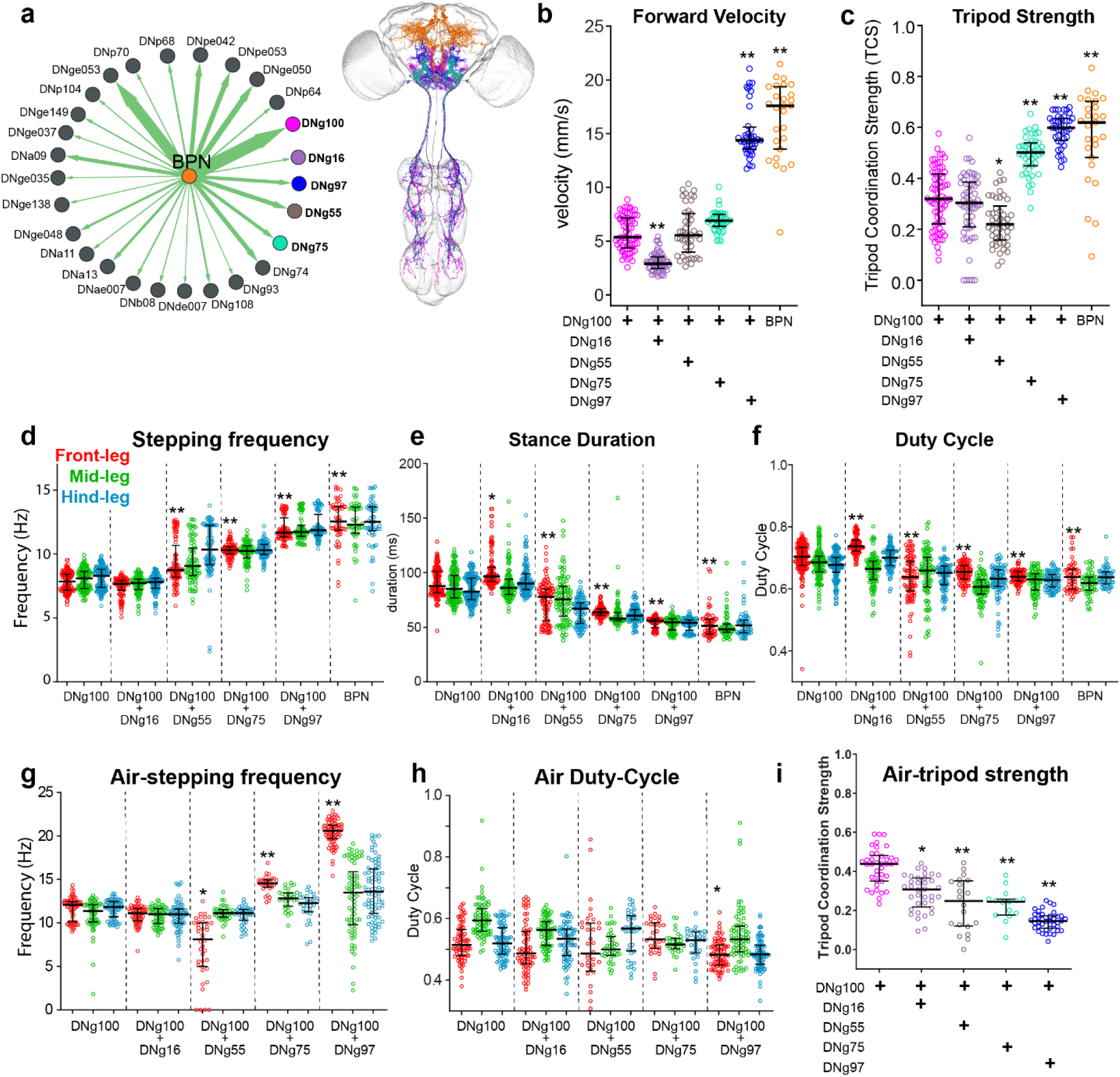
Co-activation of descending pathways modulates stepping rhythms and walking speed: **a.** Connectivity between BPN and descending neurons or DNs (left) and the EM traced anatomy (right) of BPNs and subset of DNs analyzed in following panels. Green arrows indicate excitatory connections, and red arrows indicate inhibitory connections; arrow width corresponds to synaptic count and scales from 50 to 1,163 synapses. **b-c**. Forward velocity (**b**) and tripod coordination strength (**c**) of tethered walking flies (ball-condition), where specific sets of DNs are optogenetically activated. All flies except BPN are decapitated. **d-f.** Stepping frequency (**d**), Stance duration (**e**) and Duty Cycle (**f**) of the same flies in b-c. **g-i.** Stepping frequency (**g**), Duty-cycle (**h**) and tripod coordination strength (**i**) of decapitated groups from (**c**) in the air-condition. n = 15 to 70 trials over 2-10 flies (panels d-h has data pooled from right and left side). Kruskal-Wallis test followed by Dunn’s multiple comparison with respective fly’s (**b,c,i**) or leg’s (**d-h**) DNg100 data. For d-h, showing statistical markers only for front-legs for clarity. (**p <0.001, * p< 0.05).

We first focused on within-segment left-right alternation. DNg100 stimulated flies showed strong antiphase coupling (180°) across all leg-pairs and conditions (Fig. 3d and Extended Data Fig. 4a-c). The persistence of left-right coupling after removal of proprioceptive inputs including FeCO silencing (Extended Data Fig. 3), implicates central circuits underlying left-right alternation. We also noted that left-right coupling was stronger in front legs compared to other legs, especially in air and amputated-conditions (Extended Data Fig. 4i), implying stronger central coupling pathways for front legs alternation. DNg97 stimulated flies also showed left-right antiphase coupling across conditions (Fig. 3e and Extended Data Fig. 4a-c) only for the actively stepping front-legs. The coupling was much stronger for the air and amputated conditions (Extended Data Fig. 4a-c) underscoring relevance of front-leg central coupling circuits. MDN stimulated flies in contrast showed no left-right coupling in any of the leg pairs including the rapidly stepping hind legs in air condition (Fig. 3f and Extended Data Fig. 4a-h). These results show that while DNg100 and to some extent, DNg97 are able to recruit these left-right alternation central circuits, MDN stimulation does not. This implies a task-specific recruitment of inter-leg coupling pathways by brain commands (strong coupling during DNg100 driven forward walking versus weak coupling during MDN driven backward walking).

We next looked at across-segments coupling and found that in case of DNg100 stimulation, the adjacent ipsilateral legs (i1-i2, or i2-i3) are antiphase whereas the “tripod” related leg pairs (i1-c2, c2-i3 and i1-i3) show in-phase movements across all paradigms (Fig. 3d, Extended Data Fig. 4d-h). This implies role of central circuits in across-segments coordination. As expected, given only a subset of legs show stepping across setups for DNg97 (front legs) or MDN (hind legs), there is not much across-segments coupling in these cases, highlighting once more that the core stepping CPG is local to each leg-neuromere and could operate in a decoupled state (Extended Data Fig. 4d-h).

Further, both the phase plots and coupling strength (PLV) show that when FeCO contribution is removed either by amputation (Extended Data Fig. 4a-h) or by genetic silencing (Extended Data Fig. 3b), it slightly weakens most of the inter-leg couplings. This suggests that while FeCO inputs are not necessary for inter-leg coordination they can strengthen existing central couplings.

So far, we performed uniform manipulations to all legs of the fly. But work in cockroaches^38^ and stick insects^64–66^ shows that proprioceptive inputs from one stepping leg can impact movement of legs in adjacent segments. To address this, we activated DNg100 bilaterally in flies with different sets of legs completely removed (no sensory input from the leg) and quantified air-stepping. We found that flies without front legs showed much weaker left-right coupling in mid-legs (i2-c2) and to some extent also hind-legs (i3-c3), compared to flies with front legs intact (Extended Data Fig. 6a-b). This shows that intact front leg alternation strengthens mid-leg alternation and to a smaller extent also hind-leg alternation. In contrast, front-legs alternation (i1-c1) is only slightly weakened and remains overall much stronger than other couplings even when other legs are removed (Extended Data Fig. 6a-b). This impact of front leg stepping on mid and hind-legs, could be therefore directly driven by proprioceptive feedback or due to proprioceptive feedback impacting front leg CPGs which in turn impact other legs, or both. These results underscore our original observation that DNg100 more strongly recruits the front-leg alternation circuits compared to other segments.

These results imply that DNg100 directly recruits inter-leg coupling circuits in the nerve cord. In all our experiments so far, DNg100 is stimulated bilaterally in decapitated flies. Are bilateral DNg100 inputs necessary to recruit the inter-leg coupling pathways? To address this, we used a stochastic expression strategy (Extended Data Fig. 5 and Methods) to stimulate DNg100 axons, unilaterally in decapitated flies. Unilateral DNg100 stimulation drove robust stepping in the ipsilateral front, mid and hind legs across both conditions (Fig. 3g). We observed little to no stepping in contralateral legs in the ball-condition (Fig. 3g) leading to turning with low forward and angular velocity (Extended Data Fig. 5b). However, in the air condition, in addition to stepping in all ipsilateral legs, we also observed stepping in contralateral front and hind legs, with no stepping in contralateral mid-legs (Fig. 3g). The contralateral air-steps were atypical and in case of hind legs, often comprised of small-amplitude oscillations (Supplementary video 7), however both front and hind leg movements are at typical stepping frequencies (Fig. 3g) implying a CPG origin for these movements. We found that ipsilateral inter-leg coupling (i1-i2 and i2-i3) was preserved in these flies (Fig. 3h). The contralateral front legs stepped better than hind legs and also showed stronger left-right coupling (i1-c1) compared to hind legs (i3-c3, Fig. 3g and Extended Data Fig. 5c). Thus, only a subset of inter-leg couplings was preserved on unilateral activation of DNg100. Most notably, no i2-c2 coupling was observed since contralateral mid-legs did not step.

Taken together, these data suggest existence of central pathways for both within segment and across segment inter-leg coordination. Notably, we uncover differences in coupling strengths across different segments (i1-c1 > i3-c3 > i2-c2) and show proprioceptive inputs strengthen these couplings. Finally, we find that descending brain commands are positioned to flexibly modulate inter-leg coupling strengths by directly recruiting these central pathways in a task-specific manner.

### Co-stimulation of descending inputs modulates stepping rhythm to control walking speed

Having characterized intra-leg and inter-leg coordination of the stepping rhythm, we now wondered if this observed stepping rhythm of decapitated flies could be modulated to change the walking speed of the animals. Spontaneously walking wild-type flies explore a wide range of velocities (∼0-25 mm/s) and stepping frequencies^35–37,45,61,67^. By recording DNg100 activity in spontaneously walking flies, we previously showed that DNg100 activity correlates with and precedes forward velocity changes^46^. However, DNg100 stimulated flies rarely walked faster than 8mm/s (Fig. 1b). By titrating optogenetic light intensity, we could get DNg100 stimulated flies to walk slower (Extended Data Fig. 7a-b, similar to results in a recent study^26^). This shows that the stepping frequency could be tuned by changing the DNg100 input to the leg CPGs. However, this tuning of stepping frequency was not that apparent in the air-condition where we observed a much more variable and often a binary response (stepping at high frequency, or no stepping), suggesting that proper sensory feedback was required for slowing down the stepping rhythm. This is consistent with a previous study showing that an amputated leg-stump of spontaneously walking flies doesn’t slow down even if the intact five legs slow down^36^.

But DNg100 stimulation could never drive higher velocity walking, even if we tested “head-intact” flies or flies with double copy numbers of transgenes (Extended Data Fig. 7e). Thus, DNg100 stimulated flies have access to a small slice (∼0-8 mm/s) of the total velocity range (∼0-25 mm/s). It suggests that perhaps additional DNs need to be co-activated along with DNg100 for accessing higher velocities and higher stepping frequencies. Bolt Protocerebral Neurons (BPNs), discovered in our previous work^45^, drive walking at high velocities with strong tripod coordination and high stepping frequencies (Fig. 4a-d). Connectomics^68,69^ shows that BPNs are upstream to a population of DNs^46^ including DNg100 and DNg97 (Fig. 4a). Another recently discovered forward walking regulating neuron (RRN) converges on the BPN outputs^70^. This suggests that other DNs recruited by BPN could be working in concert with DNg100 to drive higher velocity forward walking.

We therefore decided to co-activate DNg100 with other BPN downstream DNs in decapitated flies to address nerve cord specific mechanisms for speed control. In addition to DNg97, we could generate or identify genetic drivers (Fig. 4a, Extended Data Fig. 7d) for three other DNs downstream to BPN: DNg16, DNg55 (BDN3^46^) and DNg75 (cDN1^46^). While activating DNg97 by itself induces slow forward walking in decapitated flies (Fig. 1), activating DNg16, DNg55 or DNg75 did not induce any walking in decapitated flies (Extended Data Fig. 7e). By combining the genetic drivers, we could co-activate pairs of DNs (Extended Data Fig. 7d). Co-activating DNg16 with DNg100 did not speed up the velocity or stepping frequencies (Fig. 4b) and in fact led to a small decrease. Co-activating DNg100 with DNg55 led to increased variability in walking output (Fig. 4b) and although there was a statistically significant increase in stepping frequency (Fig. 4d) there was an overall decrease in inter-leg coordination leading to a lower tripod coordination (Fig. 4c), implying overall uncoordinated stepping movements. DNg100 + DNg75 coactivated flies showed a trend towards increased walking velocities and an obvious statistically significant increase in both stepping frequency and tripod coordination, compared to DNg100 stimulated flies (Fig. 4b-d). In contrast, co-activating DNg100 with DNg97 led to a major increase in walking velocities (Fig. 4b), stepping frequencies (Fig. 4d) and tripod-coordination (Fig. 4c), all of which now almost reached BPN levels.

Stride length is another parameter that can impact walking velocities. BPN activated flies showed a slightly larger stride length than all other groups (Extended Data Fig. 7g) and this parameter is probably also controlled through other BPN downstream DNs not tested here^71^. DNg100+DNg97 stimulated flies showed a slight increase in stride length compared to DNg100 case. On the other hand, DNg100+DNg55 and DNg100+DNg75 stimulated flies showed a slightly shorter stride length compared to DNg100 stimulated flies and this could explain why walking velocity of these flies shows only a modest increase despite a bigger increase in stepping frequencies.

Overall across all genotypes, in our experimental conditions stepping frequency changes were much more robust and relevant to our goal, and hence will be our main focus here. A step-period comprises of a stance period and a swing period and stepping frequency can change when either of these periods change. We found a strong reduction in stance duration across genotypes as stepping frequency increased (Fig. 4e). Swing duration also showed a similar trend although differences in swing duration are very small (Extended Data Fig. 7f). We also found that the “duty cycle” (Stance period/Total Step Period) followed a similar trend as the stance period, indicating a dominance of stance period in driving stepping frequency changes, as has been suggested from analysis of spontaneously walking flies^37,61^ (Fig. 4f). Given that, across all the coactivation groups we can tile the entire stepping frequency range (Fig. 4d) available to walking flies, we infer that co-recruiting other DNs along with DNg100 could be a reliable strategy for modulating stepping frequencies and walking velocities.

To address how these stepping frequency changes depend on sensory feedback, we repeated the above experiments without a walking substrate, and quantified air-stepping for each of the above groups (except BPN because we cannot perform air-stepping in intact flies since these flies tend to enter flight like postures). We observed that both DNg100+DNg75 and DNg100+DNg97 stimulated flies increased stepping frequencies even without the substrate (Fig. 4g). However, in both cases, but even more pronounced for DNg100+DNg97, we observed that front legs stepped at much higher frequencies (∼22 Hz which is almost twice as fast as DNg100) compared to middle and hind legs which were stepping only slightly faster than DNg100 levels. This resembles an additive phenotype of DNg100 (all legs step at ∼11 Hz) and DNg97 (only front legs step at ∼11 Hz) and strongly suggests that these DNs could be converging on the same CPG module which now receives a summed input from the DNs. The duty cycle for all air-stepping sets, was mostly maintained around 0.5 which shows these are sinusoidal step-cycles as seen earlier for air-stepping (Fig. 2). But now, especially for DNg100+75 and DNg100+DNg97 cases, since front legs step at a different frequency than other legs, this leads to a low tripod coordination strength during air-stepping (Fig. 4i). It is remarkable that when substrate contact is intact, all legs end up stepping at similar frequencies with strong inter-leg coordination (Fig. 4c). Thus, the mechanical coupling and proprioceptive feedback, is able to overcome the centrally imposed differences in stepping rhythms leading to a coordinated high-speed walking state. In contrast to DNg100 stimulated flies where air-stepping showed stronger tripod coordination than ball condition (Fig. 3c), here substrate contact is essential to drive an even better tripod coordination (Fig. 4c). This exemplifies the tight relationship between central and peripheral inputs on inter-leg coupling.

Taken together, we show that walking velocity and stepping frequency control is a function of both sensory feedback as well as descending inputs (beyond DNg100) to the nerve cord.

### Kinematics-guided connectome search identifies neural circuit motifs underlying a coordinated six-legged walking pattern

We have shown multiple lines of evidence that implies each leg of the fly is controlled by its own independent CPG. This CPG should therefore be composed of leg-neuromere specific local interneurons. DNg100 and DNg97 drove stepping in front legs in sensory deprived conditions (air and amputated) with similar stepping frequencies and similar kinematics (Fig. 1e, Extended Fig. 2a). Their coactivation resulted in front legs air-stepping at double the frequency (Fig. 4g). We therefore hypothesize that DNg100 and DNg97 converge on the same CPG module controlling front leg stepping. To find such neurons, we looked at shared outputs of these two DNs in the front-left and front-right leg neuropils in the MANC (male adult nerve cord) connectome^8^ (Extended Data Fig. 8a and Supplementary Table 3). Of the top shared outputs across left and right hemi-cord connectomes, five are leg-neuromere specific interneurons (Extended Data Fig 8a, see Methods). We therefore focused on anatomy and connectivity of these five local interneurons. We found that they form a recurrently connected motif and this motif repeats in every leg neuromere of the fly nerve cord (Fig. 5a-b). DNg100 drives stepping in all six legs and is indeed directly upstream to this motif in all leg neuromeres (Fig. 5b). On the other hand, DNg97 predominantly connects to this motif in front leg neuromeres, with sparser connections in mid-leg and hind-leg neuromeres (Fig. 5b), reflecting the behavioral phenotype in sensory-deprived conditions (Fig. 1e). MDN which drives hind-leg stepping in sensory-deprived conditions (Fig. 1e), only connects to this motif in the hind-leg neuromere (Fig. 5b). We showed that there are kinematic differences between a forward and backward stepping hind leg (Fig. 2a-e, Supplementary Video 5) and reflecting this, there are differences in the way MDN and DNg100 connect with this motif in the hind leg neuromere. We found this motif by considering only the mono-synaptic downstream neurons to the DNs. And even though the entire CPG might also consist of other neurons beyond this motif (see Extended Data Fig. 9a for expanded CPG that considers mono- and di-synaptic outputs of DNs), we propose this five-neuron motif as a core component of the walking CPG and refer to it as the “CPG-motif” in this manuscript. To further strengthen our hypothesis, we performed an identical analysis on other more recently available connectome datasets that includes a female connectome^47,48,72^, and once more ended with the same motif (Extended Data Fig. 8b-c and see Methods for selection criteria).

**Fig. 5:**
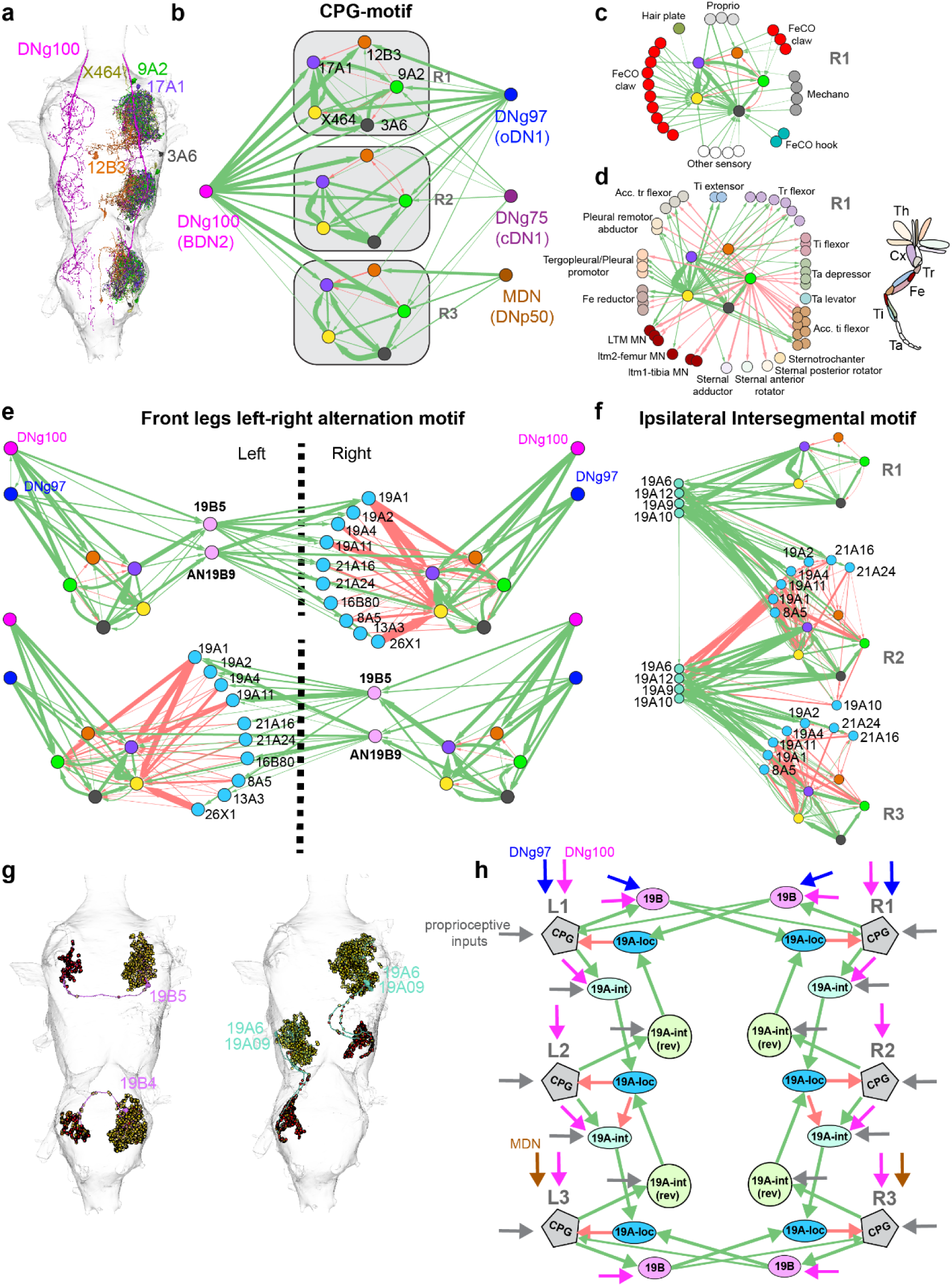
Kinematics guided connectome search isolates putative circuit motifs for intra-leg and inter-leg coordination: **a-b.** Anatomy (**a**) and connectivity (**b**) of a putative CPG-motif identified through a behavior-results guided connectome search (Extended Data Fig. 8). **c.** Direct proprioceptive inputs to CPG-motif (also see Extended Data Fig. 9). **d**. Leg motor neuron outputs of the CPG-motif (left) and corresponding leg-muscles (right, schematic adapted from Marin et al.^72^) **e.** Putative circuit motif for within-segment left-right alternation of front legs, left to right (top) and right to left (bottom). CPG-motif neurons are color-coded as in (**b**). **f.** Front to mid and mid to hind ipsilateral inter-segmental connectivity pathways. CPG-motif neurons color coded as in b. Green arrows denote predicted excitatory connections and red arrows denote predicted inhibitory connections. Arrow width is proportional to synapse number. Synapse count ranges were 5-481 (**b**), 3-479 (**c**), 5-479 (**d**), 5-479 (**e**, top), 5-546 (**e**, bottom), and 5-660 (**f**). **g.** EM traced anatomy of example 19B-commissural left-right alternation neurons for front and hind legs (left) and 19A-intersegmental neurons (right). **h.** A simplified scheme of the predicted neural circuit motifs for inter-leg coordination, conserved across three connectome datasets. Arrow thickness in (**h**) is for visualization only and does not represent synaptic weight.

This CPG-motif is also directly upstream to relevant leg motor neurons and directly downstream to proprioceptors (Fig. 5c-d and Extended Data Fig. 9b, Extended Data Fig. 10a), most notably FeCO neurons which are implicated in controlling the stepping frequency (Fig. 1e and Extended Data Fig. 1e, also ^73^). This CPG-motif is also downstream to DNg75 (Fig. 5b) that led to increased stepping frequency on co-stimulation with DNg100 (Fig. 4) Further, this CPG motif identified here, based purely on shared connectivity across behaviorally relevant DNs, shares multiple neuronal types (17A1, X464 and 3A6 from Fig. 5 and INXX466, IN19A007 from Extended Fig. 9a) with a CPG motif identified through a modeling approach (expanded motif from ^26^). This modeling study showed that such recurrently connected interneurons are capable of converting a constant DN input into a rhythmic leg-motor neuron output^26^.

We next searched for candidate left-right coupling neurons. Our experimental data shows that: 1) left-right alternation is strongest for front legs (i1-c1), 2) activating DNg100 and DNg97 drive front leg alternation, and 3) activating a single DNg100 axon unilaterally, drives contralateral front and hind legs to step but no stepping is observed in mid-legs. We therefore first searched for short commissural coupling pathways in the front leg neuropils. (a single interneuron couples left and right side, or two interneuron layers between left and right CPGs). Multiple such pathways could be initially identified in the connectome (most of which have been described in a comprehensive connectome analysis study^74^). But if we filter for first layer of interneurons that receive input from three ipsilateral sources, DNg100, DNg97 and CPG-motif, and are conserved across left and right hemi-cord, we home-in on two strongly connected commissural interneurons of the 19Bhemilineage (AN19B9 and 19B5, Fig. 5e, see Methods). These 19B commissural interneurons are predicted to be excitatory (cholinergic^8,75^) and directly upstream to a few of the contralateral CPG motif neurons (green arrows originating from 19B and connecting to CPG nodes on contralateral side in Fig. 5e, Supplementary Table 3) and could explain why we get movements of contralateral front legs with unilateral DNg100 stimulation. However, the stronger contralateral connection is through an intermediate layer of numerous local inhibitory interneurons (predominantly of the 19A hemi-lineage) that receive strong excitation from 19B commissural neurons and provide strong inhibition to the local CPG (Fig. 5e). This 2-layer contralateral inhibition motif is a strong candidate for the dominant left-right alternation in front legs, observed across all our experiments. Interestingly, this particular motif of 19B commissural neurons was absent in the mid-leg neuropil, but a similar 19B->19A motif could be identified in the hind leg neuropil (Extended Data Fig. 11a) and could explain hind-leg movements on unilateral DNg100 excitation and hind-legs left-right alternation. In case of mid-legs, we found another interneuron that formed a similar motif (Extended Data Fig. 11b) albeit with weaker connectivity that could explain weaker mid-leg left-right alternation in our behavioral data. Also, consistent with our behavioral data, MDNs are not upstream to the hind-leg commissural neurons (Extended Fig. 11a) and explains absence of strong left-right alternation in MDN stimulated flies.

To explain the ipsilateral across-segments antiphase coupling (i1-i2 and i2-i3) observed in our experiments, we searched for conserved ipsilateral intersegmental coupling interneurons, here focusing mainly on CPG to CPG connectivity. Through a similar approach (see Methods), we once more identified a 2-layer connection, where the first layer is excitatory and second layer is inhibitory (Fig. 5f). Remarkably, while this analysis was done completely independently of the contralateral coupling analysis, we found that the inhibitory layer is largely comprised of shared neurons across the two motifs (mainly 19A-local interneurons, Extended Data Fig. 11c). The excitatory layer is mostly composed of 19A-intersegmental interneurons which is a unique subset of the 19A hemi-lineage with a different neurotransmitter identity discussed in a recent connectome and lineage analysis study^72^. We also found many of the interneuron types shared across i1-i2 and i2-i3 coupling, and only show these common interneuron types in Fig 5F. We also found similar motifs in the opposite direction (hind->mid->front), which shared the same inhibitory local interneurons layer (Extended Fig. 11d). These motifs in both directions would lead to antiphase i1-i2 and i2-i3 coupling observed in all DNg100 stimulation experiments. The fact that front legs can strongly influence mid-leg coupling whereas hind-legs do not, indicates that i1->i2 coupling is more dominant in our experimental conditions compared to i3->i2 coupling. When we looked for DNg100 inputs to these 19A-intersegmental neurons, we indeed found that DNg100 strongly recruits the front->mid->hind 19A intersegmental neurons and only weakly recruits a single mid->front interneuron and none of the hind->mid interneurons.

For the sake of brevity, by just considering lineage 19 interneurons, we can explain inter-leg couplings observed across all our experiments (Fig. 5g-h). 19B-commisural->19A-local coupling is responsible for left-right alternation (strongest for front, weaker for hind and absent for mid legs). 19A-intersegmental->19A-local coupling explains ipsilateral antiphase coupling between adjacent segments. The recruitment of shared local inhibitory neurons (19A-local) by commissural and intersegmental excitatory neurons provides an efficient framework for antiphase coupling between different leg-pairs.

There are several points in this pathway where proprioceptive inputs (including hair-plates, FeCO and CS) can strengthen the couplings, both at CPGs (Fig. 5c, Extended Data Fig. 10a) as well as lineage 19 (and other identified) interneurons (Extended Data Fig. 10b). These could be sites (Fig. 5g) where stepping phase feedback re-enforces the inter-leg coordination and help with “on-the fly” adjustments to the stepping rhythm. Taken together, this kinematics-guided connectome analysis has uncovered putative neural circuit motifs underlying a coordinated six-legged walking pattern.

## Discussion

### Empirical evidence for walking CPG in *Drosophila*

By optogenetically stimulating rhythmic stepping in decapitated and deafferented preparations, we provide unprecedented evidence of leg-specific walking CPGs in *Drosophila*. This strengthens and extends claims for role of centrally generated rhythms, made in previous studies that relied on spontaneous walking *Drosophila* with either genetic silencing of proprioceptors^27^ or physical amputation of one of the six legs^36^. By amputating distal leg-joints, and in separate experiments by immobilizing proximal leg-joints, we show that each leg-joint can oscillate on its own without sensory feedback dependent on other leg-joints further underscoring central origins for observed motor rhythms. These observations imply a degree of modularity even within each leg’s CPG and is consistent with previous work in other insects^5,17^ and mammals^20,21,44^. More importantly, the specificity and reproducibility of neuronal manipulations in this study allowed us to precisely quantify walking coordination and constrain connectome search to identify putative CPG-motifs.

A recent connectome based modeling study isolated a CPG circuit motif capable of converting a constant DNg100 input into a phasic leg motor neurons output^26^. It is reassuring that the CPG motif we identified based on our empirical data shares several neurons (IN17A001, IN3A006, INXX464 from main CPG-motif and INXX466, IN19A007 from expanded CPG-motif) with CPG candidates identified by Pugliese et al.^26^, through modeling work, while also presenting candidates that were not identified in the modelling study. (IN12B003 and IN9A002). We found that IN12B003 identified in our work, but absent from Pugliese et. al CPG-motif, has a strong influence on tibia-flexors, a walking-relevant group of motor neurons that did not show rhythmic activity in the modeling work. We therefore speculate that the actual CPG might comprise neurons beyond what are predicted in either of these studies. Nevertheless, these putative CPG motifs (from our work and the modeling study^26^) and the experimental assays established here, provide perfect entry points into functional characterization of the CPG neurons in future. We also noticed that some of the CPG-motif neurons identified in our work (see expanded motif in Extended Data Fig. 9a) are shared with predicted grooming CPG neurons^76^. Our core CPG-motif also receives direct inputs from some of the grooming command pathways^77,78^ and another leg movement-driving DN (DNb08) (Extended Data Fig. 12 and Supplementary Table 3). Given the modularity of the CPG-motifs suggested by our experimental data, it is conceivable that different motor rhythms like walking and grooming employ party overlapping CPG-motifs.

Our work goes beyond identifying individual leg CPGs and provides evidence for central and peripheral pathways relevant for inter-leg coordination. Previous connectome analysis work, laid out potential pathways for inter-leg coordination, purely based on connectivity data^74^. By applying constraints from our empirical data to the connectome data and by looking at conserved pathways across connectome datasets, we could home-in on a few strong candidate pathways for left-right and inter-segmental coordination which mainly comprised of lineage-19 neurons. We can now harness the genetic toolkit of *Drosophila* and perturb these neurons in our established assays to validate their functional role in inter-leg coordination.

### Convergence with cord-circuits and mechanisms for walking across other insects and mammals

#### Descending inputs to cord-circuits for walking

We found that excitatory DNs (DNg100, DNg97 and MDN) directly recruit the pattern generating modules in each leg neuromere of the fly nerve cord. This organization is similar to excitatory reticulospinal neurons (specifically the glutamatergic LPGi/CVL reticulospinal neurons) in the mouse that recruit distributed rhythmogenic modules in the spinal cord^42^. We found that unilateral activation of DNg100 was able to recruit stepping in both ipsilateral and contralateral front and hind legs and identified potential commissural interneurons mediating this. Similarly, unilateral stimulation of LPGi/CVL reticulospinal neurons was shown to recruit ipsi- and contra-lateral rhythms in an ex-vivo preparation of the mouse spinal cord^42^.

#### CPG neurons

Many of the candidate CPG neurons identified in this work morphologically resemble CPG neurons characterized in other insects. For example, the IN12B003 inhibitory neuron identified in our work resembles a non-spiking inhibitory neuronal type (“neuron-i4” in stick insect^19^ or “neuron-i” in cockroach^16^), described as rhythmic neurons whose depolarization can reset a fictive walking rhythm. All these neurons IN12B003:fly, i4:stick-insect^19^ and i:cockroach^16^, have similar morphological and neurotransmitter characteristics: soma in the ventral posterior part of the neuropil that sends a primary neurite into the contralateral hemi-cord of the same segment where it has most of its inputs and outputs. Very few other neurons in the fly connectome have this morphology and similar ones are all from this same 12B hemi-lineage.

#### Left-right coordination neurons

Specific neurons for left-right alternation have not been identified in other insects. However, the candidates for commissural pathways (19B-commisural neurons) proposed in this work (Fig. 5g) are similar to the V0v pathway for alternation at high speeds identified in the mouse spinal-cord^79–82^ and the zebrafish spinal cord^3,83,84^. Just like the V0v pathway in vertebrates^3,44^, the identified 19B commissural neurons are excitatory and recruit populations of local inhibitory interneurons to inhibit the contralateral CPG module.

### Flexible recruitment of inter-leg coordination by descending brain commands

We found that the 19B commissural neurons and 19A-intersegmental neurons (Fig. 5h), receive direct input from walking promotion DNs suggesting that brain-commands can directly recruit inter-leg coordination pathways. Even though we found that the CPG motif is also positioned to recruit these inter-leg coordination neurons, we speculate that additional inputs from DNs are required for a stronger central coupling. A corollary of this would be that there are other DNs with inputs to the CPGs (like MDN) that bypass these commissural pathways and therefore do not impose strong coupling constraints. Why would an animal benefit from such an arrangement where inter-leg coupling could be strengthened or weakened as needed by descending brain commands? Stick insects walk slowly on precarious terrains (like sticks) where they benefit from constantly updating their inter-leg coordination based on sensory feedback from the legs^5,66^. On the other hand, cockroaches run at high-speeds on flat predictable terrain (like a bathroom floor) and rely on strong central coupling^28,39^. We previously showed that DNg100 activity is strongly correlated with forward walking velocity of the fly, whereas MDN is recruited during slow backward walking^49,85^ where the animal moves in a direction opposite to most of its high acuity senses (slow precarious walking conditions like the “stick insect” case). We hypothesize that DNg100 directly recruits inter-leg coordination motifs to scale the central couplings as a function of forward velocity (stronger coupling at higher velocity), whereas MDN does not recruit these pathways and rather relies on sensory feedback for inter-leg coordination during slow backward walking. This provides a coherent framework for flexibly scaling central versus peripheral contributions to inter-leg coordination in a task-specific manner.

### An analog of a front-wheel drive for acceleration in *Drosophila*

We found that co-stimulating additional DNs along with DNg100 led to higher walking speeds. Specifically, DNg100+DNg97 co-activation led to very high-speed walking and by comparing walking on the ball to walking in air, we showed that this increase is primarily driven by an increased front-leg stepping drive. This has some interesting implications: 1) Convergent outputs of DNg100 and DNg97 in the front-leg neuropil connectome suggest that it is likely the same CPG-module that is now oscillating at double the frequency during DNg97+DNg100 co-stimulation as compared to individual DN stimulation. This suggests a different mechanism for speed control compared to findings in zebrafish that show distinct premotor modules recruited at different speed ranges^3,86^. However, we do not believe DNg97+DNg100 co-stimulation is the only way to increase walking speed and other DNs (including DNg75 in this work) could recruit additional interneuron populations to increase speed in a similar manner as described in the zebrafish case. 2) It also shows that while stepping frequency was now different across segments during “air-stepping” leading to weak inter-leg coordination, the sensory feedback and mechanical coupling in the ball-condition now enforced strong inter-leg coordination. Indeed, in the connectome we found strong proprioceptive inputs to the 19A-intersegmental interneurons (Fig. 5g, Extended Data Fig. 10b) that could be responsible for this effect. A previous study showed importance of sensory feedback in inter-leg coordination at slower walking speeds^36^. Our results now show that sensory feedback is important for inter-leg coordination even at the very high end of the speed spectrum. 3) It shows a unique mechanism for rapid acceleration akin to a “front-wheel-drive (FWD)” mechanism for automobiles. Across animals ranging from cockroaches^87^ to cheetahs^88,89^, hind legs have been shown to be the main drivers of propulsion and acceleration (RWD or Read-Wheel-Drive analogous). We speculate that in *Drosophila*, there would be other DNs that also increase stepping frequencies specifically of hind-legs, or all-legs. Hind legs in *Drosophila* and most other animals are typically closer to the center of mass of the animal^90–92^ and also provide stronger propulsive forces making them an ideal choice for sustained high speed locomotion. Front legs on the other hand could be more relevant for directional control. We speculate that this FWD mechanism discovered here, would be primarily used for transiently accelerating in a directed manner under specific behavioral contexts. Consistent with this hypothesis, DNg97 neurons involved in the FWD mechanism are one of the main outputs of DNp09 involved in object directed pursuit^45^ and also make collateral connections with DNg100 in the brain, suggesting potential of DNg97+DNg100 co-recruitment in specific contexts. It should be noted that without an approach to test mechanically decoupled and sensory-deprived stepping, it would be difficult to isolate contributions of individual legs to increase in walking speed. This FWD mechanism could therefore be relevant in other species, just not detected. Moreover, co-recruitment of DNs with DNg100 provides an opportune model to test how animals could flexibly engage in different speed control mechanisms (e.g. FWD versus RWD) in a context specific manner.

## Methods

### Experimental Animals

Experiments were performed with *Drosophila melanogaster* raised on standard cornmeal agar supplemented with baker’s yeast at 25 °C, 60% humidity, and a 12 h light-dark cycle unless stated otherwise. For optogenetic experiments, flies were collected on retinal food (400 μM all-trans retinal; Sigma-Aldrich, R2500) after eclosion and transferred to fresh retinal food 1 day before testing, and maintained in darkness throughout their life cycle. Male flies were used for all experiments unless otherwise indicated. All full experimental genotypes with exact sample size per genotype, age and source of the genetic reagents are described in Supplementary Tables 1 and 2.

### Behavioral recording and experimental setup

Experiments were performed in a custom behavioral recording setup adapted from our previous work^46^. The hardware configuration for behavioral acquisition, including camera system, calibration, and tracking pipeline, remained unchanged from the previous study. In this work, we extended the behavioral paradigm by introducing multiple testing conditions designed to systematically reduce sensory feedback while maintaining consistent optogenetic stimulation.

### Optogenetic activation in decapitated flies

Flies expressing CsChrimson^50^ in DNg100 (BDN2), DNg97 (oDN1), or MDN were cold-anesthetized and decapitated using fine micro-scissors (Vannas spring scissors, straight, 3.5 in, 2 × 0.05 mm tip; Roboz Surgical Instrument Co.). The neck was sealed with ultraviolet-cured glue (Bondic). Flies were then tethered to a 34-gauge steel pin using ultraviolet-cured glue (LOCTITE AA 3951, Henkel) and allowed to recover before testing. Behavioral responses were assessed under four conditions as follows:

#### Condition-1

##### Ball (air-supported spherical treadmill)

Following recovery, flies were placed on an air-supported spherical treadmill^46^. Only flies that maintained stable contact with all six legs and exhibited proper grip on the ball were used for behavioral experiments. To minimize variability in movement due to experimental conditions, the air stream used to rotate the ball was checked before each experiment to ensure stable rotation without excessive wobble. Additionally, flies were positioned at approximately similar distances relative to the ball, and the fiber position for optogenetic stimulation was maintained consistently across experiments. Ball motion was recorded at 50 Hz using two orthogonally positioned custom optical motion sensors, allowing quantification of forward and angular velocity.

#### Condition 2

##### Slippery surface (SS)

To test the effect of reduced mechanical coupling, the air stream was switched off and the spherical treadmill was replaced with a custom 3D-printed mount holding a 22 mm siliconized glass coverslip (Hampton Research). The polished, hydrophobic surface provided a low-traction substrate. Tethered flies were lowered onto the coverslip, allowing leg movements without effective grip while maintaining identical optogenetic stimulation conditions.

#### Condition 3

##### Air-suspended (Air)

For air-suspended trials, flies were first placed on the spherical treadmill to allow the legs to adopt a natural resting posture and avoid the hyperextension commonly observed when flies are directly suspended in air. While on the ball, flies were subjected to a single trial of optogenetic activation. The air stream was then switched off, the ball was removed, and flies were suspended in air with all six legs freely elevated ready to be tested. Flies that exhibited sustained overextension of all legs or flight-like behavior were discarded and not used for behavioral analysis. However, flies that showed clear, visible oscillations in at least a subset of trials were retained, and were further filtered on a case-by-case basis during analysis (see below).

#### Condition 4

##### Air-suspended with leg amputation (Amputated)

For maximal reduction of sensory input, flies were cold-anesthetized and decapitated, and all six legs were amputated at the mid-femur using fine micro-scissors immediately prior to tethering. Flies were inspected to ensure viability and absence of excessive damage. Amputated flies were then tethered and tested in the air-suspended condition. Given that these flies were 12-14 days old, the success rate for this preparation was low, as many flies failed to recover or did not exhibit any observable movement following this manipulation. If flies showed viable movements, a set of 10 optogenetic stimulation trials were acquired and these were further filtered during analyses (see below).

### Femoral Chordotonal Organ (FeCO) silencing during DNg100 activation

To test the effect of FeCO silencing during DNg100 activation (Extended Data Fig. 1d-e), female flies expressing CsChrimson in BDN2 neurons and GtACR1^55^ in FeCO neurons were generated using GAL4/UAS and LexA/LexAop intersectional strategies (genotypes listed in Supplementary Table 2). Use of female flies was necessary due to transgenes inserted on X-chromosome. Female DNg100 stimulated flies showed similar phenotype to male flies (Extended Data Fig. 1d-e) and suggests no sexual dimorphism in this circuit. Flies were decapitated and tethered as described above. Simultaneous activation and silencing were delivered using two optic fibers. A red-light fiber (630 nm, 66 Hz, ∼0.040 mW.mm⁻²), positioned as in all other DNg100 activation experiments, was used to activate CsChrimson, while a green-light fiber (530 nm, Thorlabs M530F2, 66 Hz, ∼0.028 mW.mm^−2^) was positioned lower and directed toward the ventral nerve cord to activate GtACR1. Flies were tested for up to 10 trials each under the same stimulation timing described below.

### Unilateral activation experiments in decapitated flies

To carry out stochastic unilateral activation of DNg100 (Fig. 3, Extended Data Fig. 5 and Supplementary Video 7), flies carrying 20XUAS-FRT>dSTOP-FRT>CsChrimson-mVenus^93^; hsFlp2 were crossed to the DNg100 split-GAL4 line (VT058557-p65ADZp; R85F12-ZpGDBD). Crosses were maintained at 19 °C, and larvae were heat shocked at 37 °C for 1.5–2 h during development to induce Flp-mediated excision of the STOP cassette, resulting in stochastic CsChrimson expression in BDN2 neurons. Adult flies were aged for 14–16 days, then prepared and tethered as described above. Unilateral DNg100 stimulation effects were assayed under ball and air-suspended conditions using the optogenetic stimulation protocol described below (66 Hz, up to ten trials per fly). Following behavioral testing, flies were dissected to determine expression patterns. Based on immunohistochemistry results (Extended Data Fig. 5a), that correlated perfectly with forward and angular velocity measurements (Extended Data Fig. 5b), flies were classified as left, right, bilateral DNg100 expression, or negative (no DNg100 expression). The unilateral groups (left and right) were used for leg-kinematics analysis.

### Segment-specific leg removal experiments

To further examine the contribution of individual leg groups and inter-leg coupling, targeted leg-removal experiments were performed in decapitated, tethered flies (Extended Data Fig. 6). Legs were amputated as close to the body as possible, and any remaining stumps were sealed with ultraviolet-cured glue to prevent movement and residual sensory input. Flies were tested in multiple configurations, including front + mid legs (hind legs removed), mid legs only (front and hind legs removed), front + hind legs (mid legs removed), front legs only (mid and hind legs removed), hind + mid legs (front legs removed), and hind legs only (front and mid legs removed), allowing comparison of motor output across different remaining leg combinations under identical stimulation conditions.

### Proximal joint immobilization

To assess the contribution of proximal hair-plate sensors (Extended Data Fig. 1f-h and Supplementary Video 4), flies were cold-anesthetized and decapitated, after which the proximal joints of the forelegs were immobilized using ultraviolet-cured glue applied up to the femur-tibia (FeTi) joint (Extended Data Fig. 1f-h). Mid and hind legs were then amputated as close to the body as possible, and the remaining stumps were sealed with glue to eliminate any residual movement. Flies were subsequently tethered and tested in the air-suspended condition to examine movement at distal joints without any feedback from proximal movements of front legs. This was done in tandem with leg-removal experiments and the intact front-legs movement data from leg-removal experiments was used as control condition here.

### Co-activation experiments in decapitated flies

Co-activation of BDN2 with other descending neurons was used to assess its influence on motor output (Fig. 5 and Extended Data Fig. 7). Flies expressing CsChrimson in combinations of BDN2 (DNg100) with DNg16, BDN3 (DNg55), cDN1 (DNg75), or oDN1 (DNg97) were tested. A new BDN2 (DNg100) stimulated dataset was generated as control for this experiment. Flies were prepared and tethered as described above. Behavioral responses were assessed under two conditions: the spherical treadmill (ball) and the air-suspended condition. Optogenetic stimulation was delivered using the protocol described below (66 Hz, up to ten trials per fly). Recording conditions were maintained consistently across genotypes and behavioral conditions to enable direct comparison of motor outputs.

### Optogenetic stimulation protocol

The optogenetic stimulation protocol was identical across all conditions above. Each fly was tested for a maximum duration of 20 min, during which up to ten trials (7 s each) were delivered. Each trial consisted of 2 s of light OFF followed by 3 s of red light stimulation (66 Hz, ∼0.040 mW mm⁻²), delivered through an LED-coupled optic fiber (625 nm; Thorlabs M625F2).

### Intensity-sampling activation in decapitated BDN2 flies

To examine how motor output scales with descending drive strength, flies expressing CsChrimson in BDN2 were tested under varying light intensities (Extended Data Fig. 7a-c). The experimental preparation and tethering were identical to those described above. Stimulation timing remained fixed (2 s OFF + 3 s ON at 66 Hz), while only the peak light power during the 3 s stimulation period was varied. The seven irradiance levels used were 0.002 0.004, 0.008, 0.016, 0.024, 0.032 and 0.040 mm⁻² (corresponding to intensity levels 1–7, respectively, in Extended Data Fig. 7a–c), measured at the fiber tip using a photodiode power meter (Thorlabs PM100D) and sensor (Thorlabs S121C) at 66Hz. Intensity levels were randomized across trials to prevent order-dependent adaptation.

### Kinematic analyses

#### 1. 2D keypoint tracking and 3D reconstruction

All behavioral data collected as were processed using an updated version of our previously published kinematic analysis pipleline^46^. Using DeepLabCut^94^ (v.2.3.11) we trained separate ResNet152 networks to track 33 points of interest on the fly body: the notum, two wing hinges and five joints per leg (thorax-coxa, coxa-trocanter, Fe–Ti, tibia-tarsus and the tarsal tip). Each sensory deprivation condition had its own dedicated networks trained with ∼6000 manually annotated training frames. The networks were re-trained until the average training error dropped below 7pixels. The resulting 2D key point tracking was triangulated using Anipose^95^ to obtain the 3D reconstruction of the fly. The average reprojection error across all tracked keypoints had to be below 20 pixels for a given reconstruction to be used for downstream analyses. Cases where the reprojection error exceeded this limit were proofread using our custom proofreading UI (see below) to arrive at better reconstructions. Across conditions, the tarsal tip tracking was prone to most error and was hence excluded from the final 3D reconstructions.

Six joint angles were calculated per leg using the framework provided by Anipose as follows:

a. <u>Thorax-coxa flexion</u>: flexion-extension angle between an adjacent ipsilateral thorax-coxa, thorax-coxa and coxa-trochanter joints of the leg of interest (calculated as the angle between the 2 vectors defined by the 3 points). The sign of this angle was chosen such that an increment represents anterior movement.
b. <u>Thorax-coxa rotation</u>: measured as the angle swept by the femur around the axis defined by the coxa.
c. <u>Thorax-coxa abduction</u>: flexion-extension angle between the contralateral thorax-coxa, thorax-coxa and coxa-trochanter joints of the leg of interest. The sign of this angle was chosen such that an increment represents lateral movement of the leg. Note that for all groups analyzed in this study this angle showed no significant range of motion and hence was omitted from the analyses.
d. <u>Coxa-trochanter flexion</u>: flexion-extension angle between thorax-coxa, coxa-trochanter and femur tibia joints.
e. <u>Coxa-trochanter rotation</u>: measured as the angle swept by the tibia around the axis defined by the femur.
f. <u>Femur-tibia flexion</u>: flexion-extension angle between the coxa-trochanter, femur-tibia and tibia-tarsus joints.

#### 2. Proofreading with custom proofreading UI (3D-PoseProof)

We developed a custom graphical user interface “3D-PoseProof” to detect tracking errors from large movie datasets and perform targeted proofreading of keypoints from multiple camera views. Given movies and 2D keypoint tracking from multiple cameras and corresponding 3D trajectories and joint angles, we detect error frames in one of two ways:

a. a sudden, atypical jump in any flexion joint angle is detected as an error in any of the constituent joints.
b. on comparing 2D trajectories of any single keypoint across 2 different camera views, any deviation in the rate of change of coordinates suggests that there is a tracking error in at least one of them.

The user can switch between these 2 error detection modes and refine the error thresholds as needed. 3D-PoseProof provides an intuitive interface where the user can easily navigate between all the detected errors, view all neighboring frames around an error, proofread 2D tracking and toggle camera views of the same frame to ensure proper tracking from all cameras. The UI is also integrated with the folder structure required by Anipose so that on completion of proofreading, the folder structure is updated such that Anipose can be readily re-run on the proofread 2D tracking.

#### 3. Post-processing, step-cycle predictions

The 3D reconstruction output from anipose is rendered in an arbitrary coordinate system. We defined a new set of basis vectors and performed a change of basis operation on these coordinates such that the new x,y and z axes were aligned along the medio-lateral, antero-posterior and dorso-ventral body axes of the fly. Given these new coordiantes, the peaks and troughs of the y-axis trajectory of the femur-tibia joint correspond to the anterior-extreme position (AEP) and posterior extreme position (PEP) of the leg, respectively (Extended Data Fig. 1b). We custom defined thresholds in the scipy find_peaks function (using prominence, distance and width thresholds) for each genotype in every experimental condition such that the peaks and troughs picked out corresponded to real step-cycles discernable on manual inspection of the movies. A single threshold was used per genotype per condition to maintain a consistent definition for a given group, but different definitions were adopted across genotypes and conditions to accommodate different kinds of steps performed (Extended Data Fig. 1c). We also crosschecked the validity of step cycles defined in this way by comparing these step cycles with those predicted using our previously published ball-fitting strategy for flies walking on the spherical treadmill (Extended Data Fig. 1b). From this comparison, we found that while for DNg100 and DNg97 stimulated flies, peaks along the AEP-PEP axis corresponded best with real step cycle transitions, peaks along the dorso-ventral axis (DEP-VEP) was the best approximation for the hind legs of MDN stimulated flies. Hence, these definitions were adopted across conditions for the respective genotypes for all analyses.

#### 4. Data quality control

All collected data was quality controlled to ensure analysis of properly tracked representative data. For all bilateral stimulation experiments, any fly showing stepping only on one body side was excluded from analysis.

##### Ball-condition

Only flies that showed a significant increment in velocity (>2mm/s) on optogenetic stimulation were retained for DNg100, DNg97 and MDN stimulated conditions.

##### Air and amputated-conditions

Once a large enough dataset was collected (>10 flies) for each genotype, each leg segment was scored as stepping or non-stepping. Based on this, the actively stepping legs were identified as the ones showing discernable oscillations in >90% of flies imaged (defined as >1Hz stepping frequency). Once the stepping legs were identified per condition per genotype, only those trials with the best step quality was retained (defined as clean step detections using a generalizable threshold across all flies and trials in the dataset).

<u>For unilateral activation</u> of DNg100 only flies which showed active stepping on all legs ipsilateral to the DNg100 axon stimulated in both ball and air conditions were retained in the analyses. <u>For segment specific leg removal experiments</u>, filtering conditions similar to that applied for the regular air suspended condition were used, where all flies showing active movement on both body sides were analyzed.

Optogenetic stimulation phenotypes tend to decline over extended stimulation. To focus on the most robust part of optogenetic stimulation, for all experiments involving DNg100 or DNg97 stimulation, only the first second of optogenetic stimulation was analyzed. For MDN stimulation experiments, a window of 1.5 seconds starting at 0.5 s post onset of stimulation was analyzed to accommodate for a slight delay in stepping onset as well as relatively slower stepping in this genotype.

#### 5. Kinematic parameter definitions

1. <u>Stepping frequency</u>: Defined as the per trial median of all inter-step intervals (inter-AEP intervals for forward stepping and inter-PEP intervals for backward stepping) over the analyzed period of the trial. If no peaks were detected in a trial, a default value of zero was assigned. For unilateral activation experiments (Fig. 3g-h) the reported value is the median over all inter-step intervals over all trials per fly.
2. <u>Average step calculation</u>: For a forward step, a step cycle was defined as the period between 2 consecutive AEPs. For a backward step, a step cycle was defined as the period between 2 consecutive PEPs. For each genotype in each condition of sensory deprivation, the time series of each joint angle from 2 consecutive step cycles was extracted and each was up-sampled to 200 frames (1s). This time normalized dataset was z-scored and averaged to get the average temporal structure (Fig. 2b-e and Extended Data Fig. 2a-b).
3. <u>Duty cycle</u>: Defined as the per trial median of the instantaneous duty cycle calculated as stance duration/ total step cycle duration (Fig. 2f-h).
4. <u>Tripod coordination strength</u>: Calculated as previously reported^37^. For each tripod set of legs (L1-R2-L3 and R1-L2-R3), the instantaneous TCS was defined as total overlapping swing duration/ (onset of the last swing – onset of the first swing). The reported value is the per trial median over all instantaneous TCSs per trial (Fig. 3c). Every step that was not part of a ‘tripod event’ was assigned a default TCI of zero.
5. <u>Inter leg phase plots</u>: For a given intersegmental leg pair, one leg was chosen as the reference (for a left-right pair this was always the left leg and for an ipsilateral pair, the more anterior segment). Given a pair of adjacent touchdown (TD) events in the reference leg, if there is at least one TD event in between these in the test leg, the phase offset between the two is calculated as 2^nd^ TD in reference – TD in test / 2^nd^ TD in reference – 1^st^ TD in reference. If there is no TD in the test leg in between 2 in the reference, no score is assigned. If there are multiple events, multiple scores relative to the reference pair are assigned independently.
6. <u>Intra leg phase plots</u>: Peaks of all joint angles in a leg were picked using scipy find_peaks function (using prominence, distance and width thresholds). Given 2 TD events in a leg, the phase offset of each joint angle to the TD event was calculated as the offset of the peak any given joint angle with respect to the first TD event of the pair (Extended Data Fig. 2d).
7. <u>PLV (Phase Locking Value)</u>: Calculated as previously defined^63^. The time series of leg oscillations along the AEP-PEP axis of the desired pair of legs, were band pass filtered (4-15Hz) with a 5^th^ order Butterworth filter. The mean absolute phase offset between 2 such signals was calculated as the difference between the instantaneous phase angles extracted from the Hilbert transform of the respective signals (Extended Data Figs. 3b and 4a-i).
8. <u>Stance duration, swing duration</u>: For a forward stepping leg, stance duration is defined as the time between PEP and immediately preceding AEP, and swing duration is defined as the time between AEP and the immediately preceding PEP. For a backward stepping leg, the definitions are consistent, except the AEP and PEP points are reversed.
9. <u>Stride length</u>: We calculate this parameter only for flies mounted on a spherical treadmill. We use our previously published^46^ ball fitting algorithm to estimate the location of the treadmill from the tarsal tip positions of the legs. Once the spherical surface is estimated, we identify time points when the legs are on the surface (stance) and off the surface (swing). The stance distance is calculated as the displacement of the tarsal tip during stance. The value reported is the per trial median (Extended Data Fig. 7g).
10. <u>Body length normalization of ball velocity</u>: Body length of each fly was measured as the distance in pixels between the front and hind thorax-coxa joints, from the 2D camera view that looks at the side view of the fly (focal plane parallel to the long body axis of the fly). Mean ball velocity during optogenetic stimulation was divided by this body length in pixels for every fly to get a body-length normalized forward velocity.

### Connectome analysis

#### 1. Finding putative ‘CPG-motif’

Male Adult Nerve Cord (MANC v1.2.3)^8^ and Male Central Nervous System (MaleCNS v0.9)^47^ connectomes were accessed using the neuprint-python package (0.5.2)^96^. Brain And Nerve Cord v626 (BANC)^48^ synapse and neuron table was downloaded from https://codex.flywire.ai/api/download?dataset=banc. We defined ‘CPG-motif’ as leg-neuromere specific interneurons shared between DNg100 and DNg97 on both sides using MANC. To find such shared neurons, top 100 outputs (neurons with synapses>=5) of DNg100 and DNg97 in T1 (front leg neuromere) were ranked based on their synaptic weights such that the neurons receiving most synapses in T1 were assigned 100th rank. Neurons whose type starts with (‘IN’,’AN’,’dP’,’EN’,‘EA’,’LB’) were retained to exclude motor neurons and descending neurons from our analysis. For each output neurons on both sides, we calculated ‘rank sum’ as sum of DNg100 and DNg97 ranks. On sorting the ‘rank sum’ in descending order, we applied a stringent cutoff of top 10 ‘rank sum’. Neurons that are shared between top 10 ‘rank sum’ of left and right sides were retained. From these lists, based on our ‘CPG-motif’ definition, we found 5 putative ‘CPG-motif’ neurons: IN03A006, INXXX464, IN12B003, IN17A001 and IN09A002 (Fig. 5a-b, Extended Data Fig. 8 and Supplementary Table 3).

These steps were repeated in MaleCNS and BANC connectome dataset and found same neurons. It is important to note that IN17A001, one of our CPG neurons, did not appear within top 10 outputs of DNg97 on the right side of VNC for maleCNS dataset. Given that they are:

1. among top 10 outputs for DNg97 and DNg100 in MANC and BANC (both left and right side),
2. among top 15 outputs of DNg97 on the right side,
3. highly interconnected with other putative CPG neurons, we decided to include this neuron in our CPG candidate list.

Note, that if we perform the same analysis without restricting to top 10 interneurons and look for conserved outputs across all connectomes and hemi-cords, we only add 3 more weakly interconnected neurons to the CPG-motif.

#### 2. Finding connectome network paths

Network paths were generated by providing source and target ids to fetch paths() from neuprint-python^96^ with synaptic threshold >=5 (we considered synaptic threshold >= 3 when sensory neurons were considered) and timeout = 60000s. These paths were later converted to a network using networkx python package(v3.6.1). Networks were generated such that each node (neuron ID) were connected to each other by edges based on their synaptic weights. To account for multiple neuron IDs of same neuron type within each leg neuromere, neurons (nodes) with same ‘instance’ were merged and weights (edge) between identical instance pairs were summed to generate ‘merged networks’. All networks were exported to cytoscape^97^ (v3.10.4) for visualization. Information on the connectivity matrices underlying the nodes shown in the figures will be made available upon request.

#### 3. Expanded CPG-motif and motor neuron influence scores

To define expanded CPG neurons, we generated a merged 2-hop network path in MANC such that T1 CPGs target front leg motor neurons via an intermediate local neuron layer. These intermediate local neurons were further filtered by retaining only neurons that receive synapses>= 250 from CPG neurons. After considering shared intermediate local neurons across left and right T1 neuromere, we identified 8 additional neurons as expanded CPG-motif. These neurons had similar connectivity between CPG and motor neurons in T2 and T3 neuromere (Extended Data Fig. 9 and Supplementary Table 3). Additionally, expanded CPG-motif connectivity with motor neurons and CPG neurons were confirmed in BANC and MaleCNS.

To categorize the influence of each CPGs onto motor neurons, ‘unsigned’ influence scores were calculated from the MANC 2-hop network from CPG to motor neurons similar to previously published method^70^. Input fractions for each neuron pair in the network were calculated as the ratio of its weights with the presynaptic partner to the total weights onto the postsynaptic neuron. Based on the input fractions, we generated a connectivity matrix (C_If_). To calculate 2-hop and 1-hop influence of CPG to motor neurons, following matrix multiplication was performed:

> Influence score matrix = C_If_ + (C_If_ * C_If_)

From the resultant influence score matrix, we considered motor neuron for each CPG neurons whose influence score was above 0.4. Motor neurons that appear in both left and right T1 network were reported.

#### 4. Three-hop contralateral network in MANC

To find candidate neurons responsible for left-right alternation in T1 (Fig. 5e), we specifically looked for a motif where activation of one side of CPGs leads to inhibition of the other. Considering that we could not find a direct inhibition between two CPGs conserved on both sides, we searched for an indirect 3-hop network. 3-hop network path from DNg100, DNg97 and CPG neurons on the left side (source) to CPG on the right (target) was searched using fetch_paths() function. On merging the network, we applied a criterion such that the source on left side innervates excitatory neurons (layer 1) which excites inhibitory neurons (layer 2) that inhibits CPGs on the right side were retained. We further considered only neurons in layer 1 and 2 based on following neuron subclasses^72^:

1. If layer 1 neurons are any of the subclass from (’BI’,’BR’,’BA’,’CI’,’CR’,’CA’) then layer 2 neurons must be ‘IR’
2. If layer 1 neurons are of ‘IR’ subclass then layer 2 must belong to any of subclasses from (’CR’,’CI’,’BR’,’BI’,’CA’,’BA’)

Similar networks were made from DNg100, DNg97 and CPG neurons on the right side (source) to CPG on the left (target). To filter for the conserved layer 1 neurons from left-right and right-left, we considered shared layer 1 neurons in T1 and their corresponding top 10 layer 2 neurons based on synaptic weights. Similar logic was applied to find layer 1 and 2 neurons for T2 and T3 (Extended Data Fig. 11a-b). However, in these two cases, we only considered paths from DNg100 and CPG while excluding ascending subclasses from the criteria.

#### 5. Three-hop intersegmental network in MANC

To find candidate neurons responsible for i1-i2 and i2-i3 alternation (Fig. 5f), we searched for a motif where activation of:

1. CPG in T1 inhibits CPGs in T2, that inhibits CPGs in T3 (Front to Hind)
2. CPG in T3 inhibits CPGs in T2, that inhibits CPGs in T1(Hind to Front).

Direct inhibition motifs were not found to be shared across left and right from T1 to T2 and T2 to T3 directions. Therefore, we searched for a 3-hop motif similar to our contralateral networks.

In case of front-to-hind network, we searched for a 3-hop network path from CPG(T1) to CPG(T2) and CPG(T2) to CPG(T3) on both sides, such that excitatory neurons (layer 1) receiving inputs from the source excite inhibitory neurons (layer 2) that outputs to the target. To filter for these neurons in a merged network, we searched for neurons based on following neuron subclasses^72^ possibilities:

1. If layer 1 neurons are of ‘II’ subclass then layer 2 neurons must be ‘IR’ or
2. If layer 1 neurons are of ‘IR’ subclass, then layer 2 neurons must be ‘II’

Following this, we searched for shared excitatory and inhibitory layer neurons between left and right side. Similar criteria were followed to create 3-hop hind-to-front network where network paths were generated from CPG(T3) to CPG(T2) and CPG(T2) to CPG(T1) (Extended Data Fig. 11d).

### Statistics

All statistical tests were performed in Matlab or Graphpad Prism. All 2-group comparisons (unless indicated otherwise) were performed using non-parametric Mann-Whitney test. All multiple-group comparisons were performed using non-parametric Kruskal-Wallis test followed by Dunn’s multiple comparison with appropriate control. Exact sample size values for each plot are reported in Supplementary Table 1.

### Code

Custom scripts were written in either Matlab 2021a or Python. A Github repository containing all code related to this manuscript will be made available post-publication on the bidaye-lab Github page (pre-publication upon reasonable request).

## Acknowledgments

We thank Michael Dübbert for technical assistance with several electronics components in the experimental setups. We thank Max Planck Florida Institute mechanical workshop for custom designed parts in all our setups. We thank John Tuthill and Bingni Brunton for sharing their CPG modeling results prior to publication. We thank Gianna Vitelli for identifying DNg55 driver line used in this study. We thank Varshitha Bojanapati and Jensen Teel for help with training and proofreading leg tracking datasets. This work was funded by Esther A. & Joseph Klingenstein Fund (SSB) and the Max Planck Society (SSB).

## Author Contributions

NS and DSK jointly led this project. NS performed all experiments and connectome analysis with support from KM, NM and SS. DSK performed all kinematics analysis with support from JP. SSB conceived the project, designed and supervised all experiments and analysis, and wrote the manuscript with support from all co-authors.

---

**Extended Data Figure 1:**
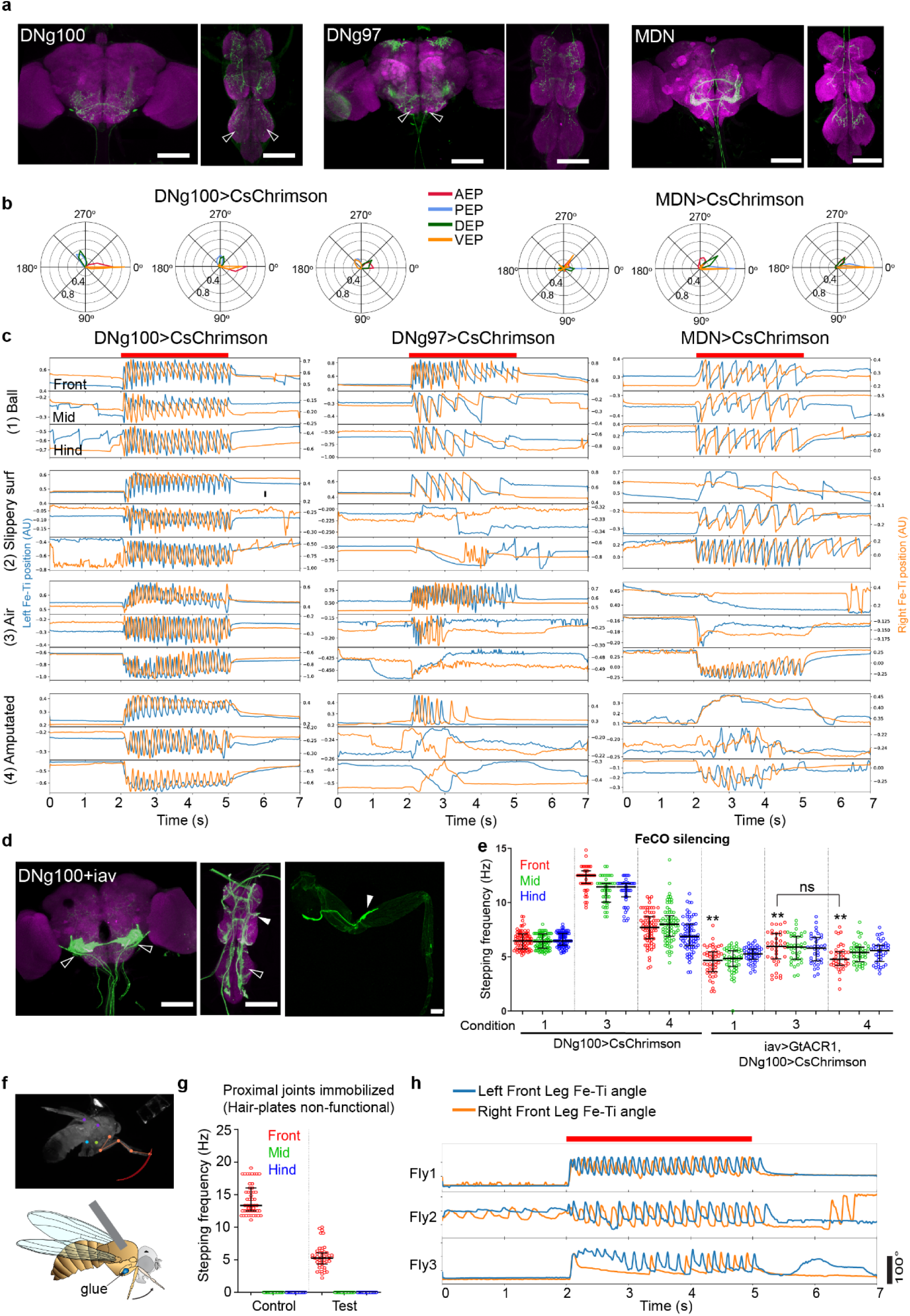
Optogenetic activation of DNs across sensory deprivation conditions drives leg oscillations. **a.** Immunohistochemistry (IHC) images of DNg100>CsChrimson-mVenus (left), DNg97>CsChrimson-mVenus (middle) and MDN>CsChrimson-mVenus lines. Brain on left and Ventral Nerve Cord (VNC) on right in each panel. Open arrowheads indicate neuron of interest. green:antiGFP and magenta:nc82 staining. scale bar = 100 µm. **b.** Circular histograms showing distribution of AEP (Anterior Extreme Position), PEP (Posterior Extreme Position), DEP (Dorsal Extreme Position) and VEP (Ventral Extreme Position) of the Fe-Ti joint position in DNg100 stimulated forward stepping legs (left) or MDN>CsChrimson stimulated backward stepping legs used to define step cycles. 0° corresponds to touch down and the event that aligns best with touch-down was used to define step cycles across conditions (AEP for all forward-stepping legs, PEP for backward stepping front and mid legs and VEP for backward stepping hind-leg). **c.** Example traces of Fe-Ti joint position for left (blue) and right (orange) front, mid and hind legs of DNg100 (left), DNg97 (middle) and MDN (right) stimulated decapitated flies across four sensory deprivation conditions from Fig. 1c. Red bar on top indicates time of optogenetic stimulation. **d.** on left, IHC image of DNg100>CsChrimson-mVenus, iav>GtACR1-YFP (nc82:magenta, antiGFP:green labels both CsChrimson and GtACR1, open arrowhead:BDN2, solid arrowhead: iav chordotonal afferents). Right, Projection of a confocal stack through prothoracic leg showing YFP fluorescence, arrow indicates the iav expression in FeCO. scale bar = 100 µm. **e.** Stepping frequency for DNg100 stimulated flies (control) and DNg100 stimulated with iav silenced flies across three conditions from Fig. 1c. 32 to 76 trials across 4 to 8 flies. **f.** Example video frame (top) with right front Fe-Ti tracking data (orange dots) during DNg100 stimulation and schematic of fly (bottom) suspended in “air” with front leg proximal joints immobilized with UV glue (top) and. **g.** Fe-Ti joint angle cycle frequency of intact front legs (control) or front legs with immobilized proximal joints (test) of DNg100>CsChrimson stimulated flies (mid and hind legs were removed for both test and control). **h.** Example trace of test flies from f. showing Fe-Ti joint angle traces during optogenetic stimulation (red bar on top) of DNg100. (also see Supplementary Videos 4). Supplementary Table 1 shows full experimental genotypes and exact sample sizes.

**Extended Data Figure 2:**
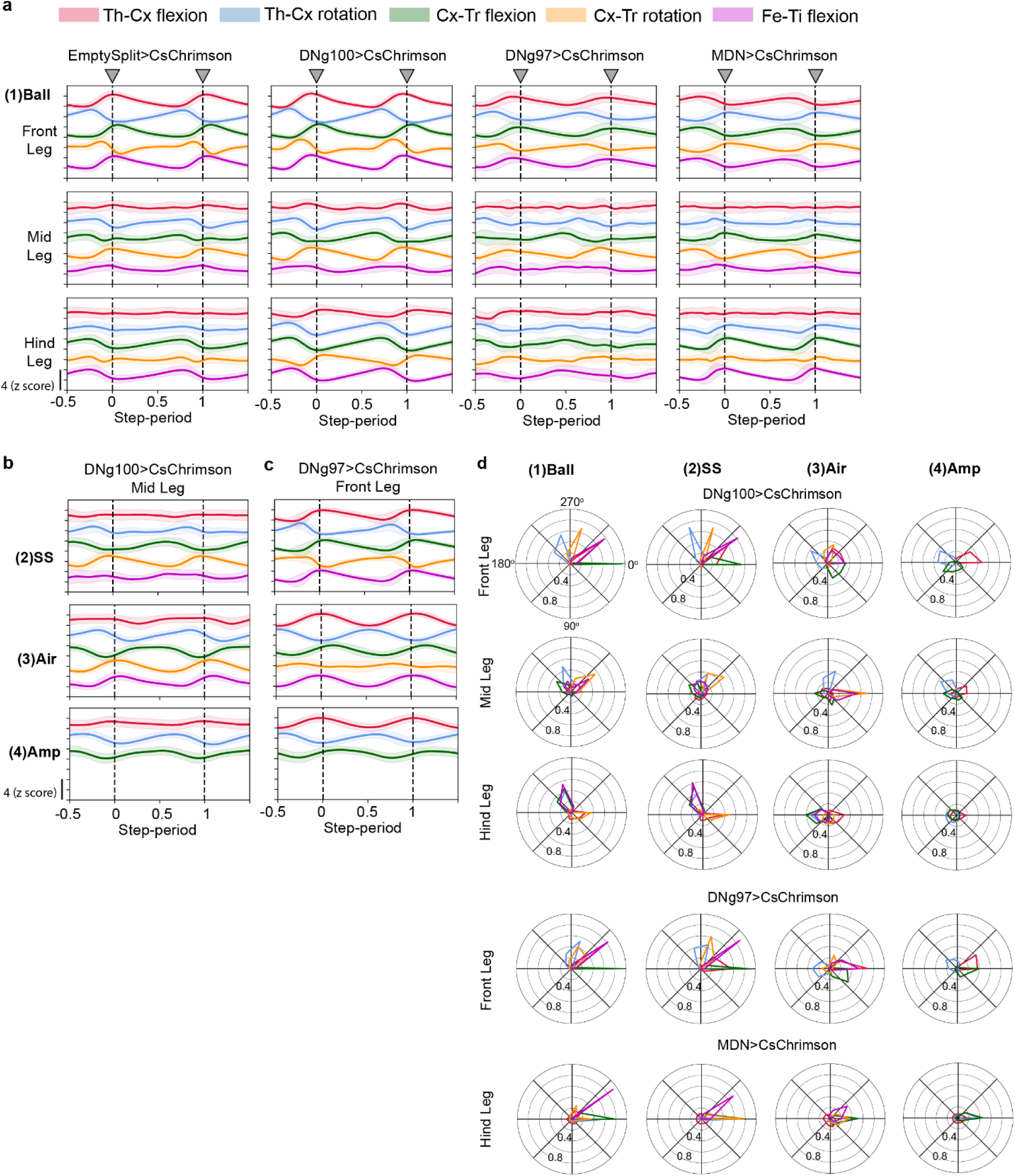
Intra-leg coordination characterization. **a.** Intra-leg joint angle traces over an averaged step cycle (mean ± s.e.m., color coded as in Fig. 2a) for control head-intact flies walking spontaneously on a ball (left) and all test flies (right three panels, decapitated flies with respective optogenetic stimulation walking on a ball). Some traces are repeated from Fig. 2b to provide comparative view across genotypes and legs. 32 – 70 trials across 6-7 flies. **b.** DNg100 stimulated fly mid leg data for corresponding panels in Fig. 2c-e. **c.** DNg97 stimulated fly front leg joint angle traces across respective conditions from Fig. 1c. **d.** Circular phase plots showing relation of each joint angle peak within a step cycle period as defined in Extended Fig 1b. Four columns represent each condition in Fig. 1c and each row represents actively stepping leg of each DN stimulated decapitated walking flies. (all legs for DNg100, only front legs for DNg97 and only hind legs for MDN, left and right leg data is pooled for all plots in this figure).

**Extended Data Figure 3:**
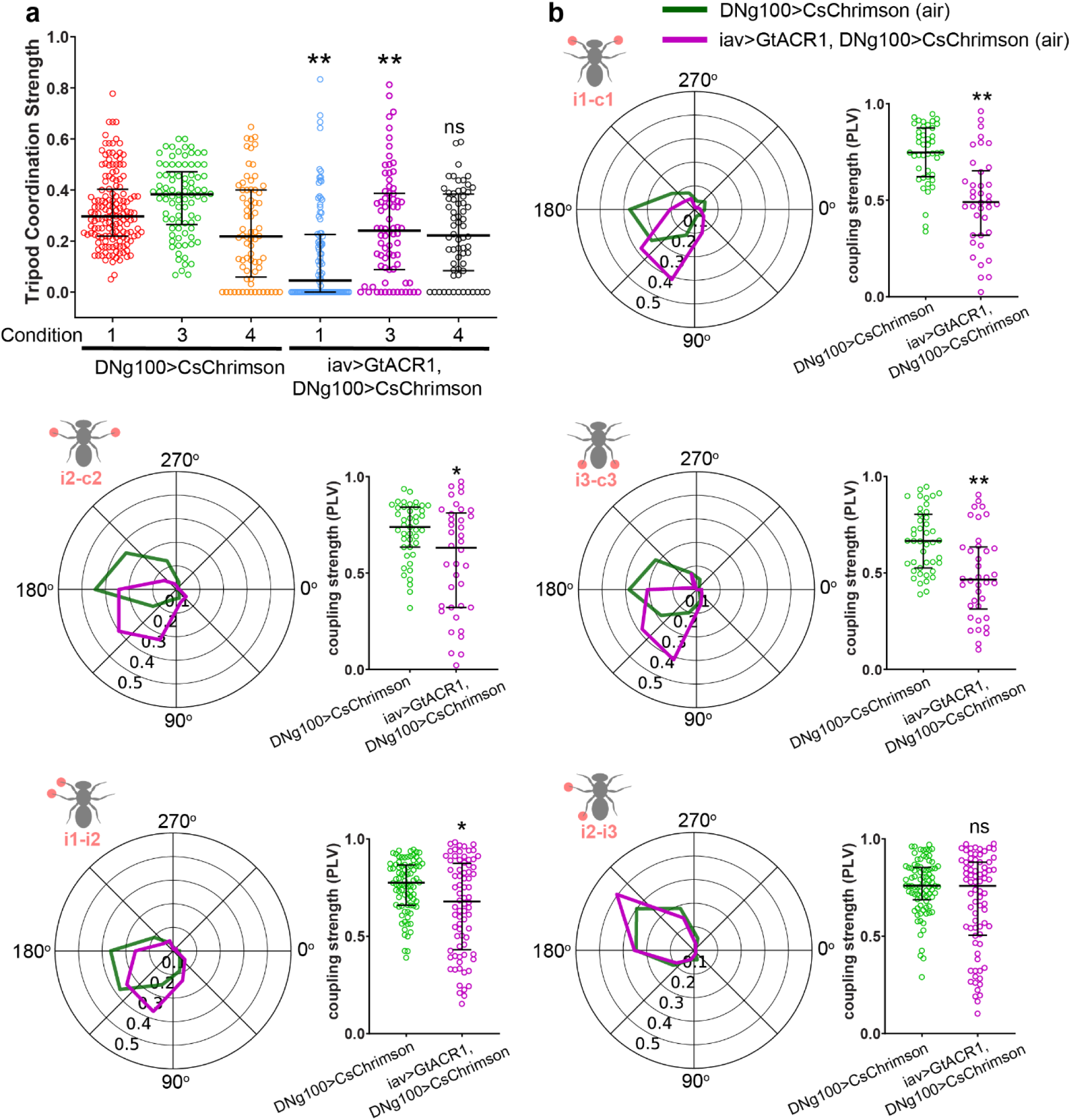
Inter-leg coordination for DNg100 activation with FeCO silencing. **a.** Tripod Coordination Strength (TCS) for control and test group across ball, air and amputated conditions. Statistics show Mann-Whitney test between control and test group for each condition (*p<0.05). 32-76 trials across 4-8 flies. **b.** Inter-leg coordination phase plots as in Fig. 3 (n= 125-749 phases), and corresponding coupling strengths, quantified as Phase Locked Value (PLV, see Methods). Statistics show Mann-Whitney test between control and test group for coupling strengths in each condition (*p<0.05, **p<0.001).

**Extended Data Figure 4:**
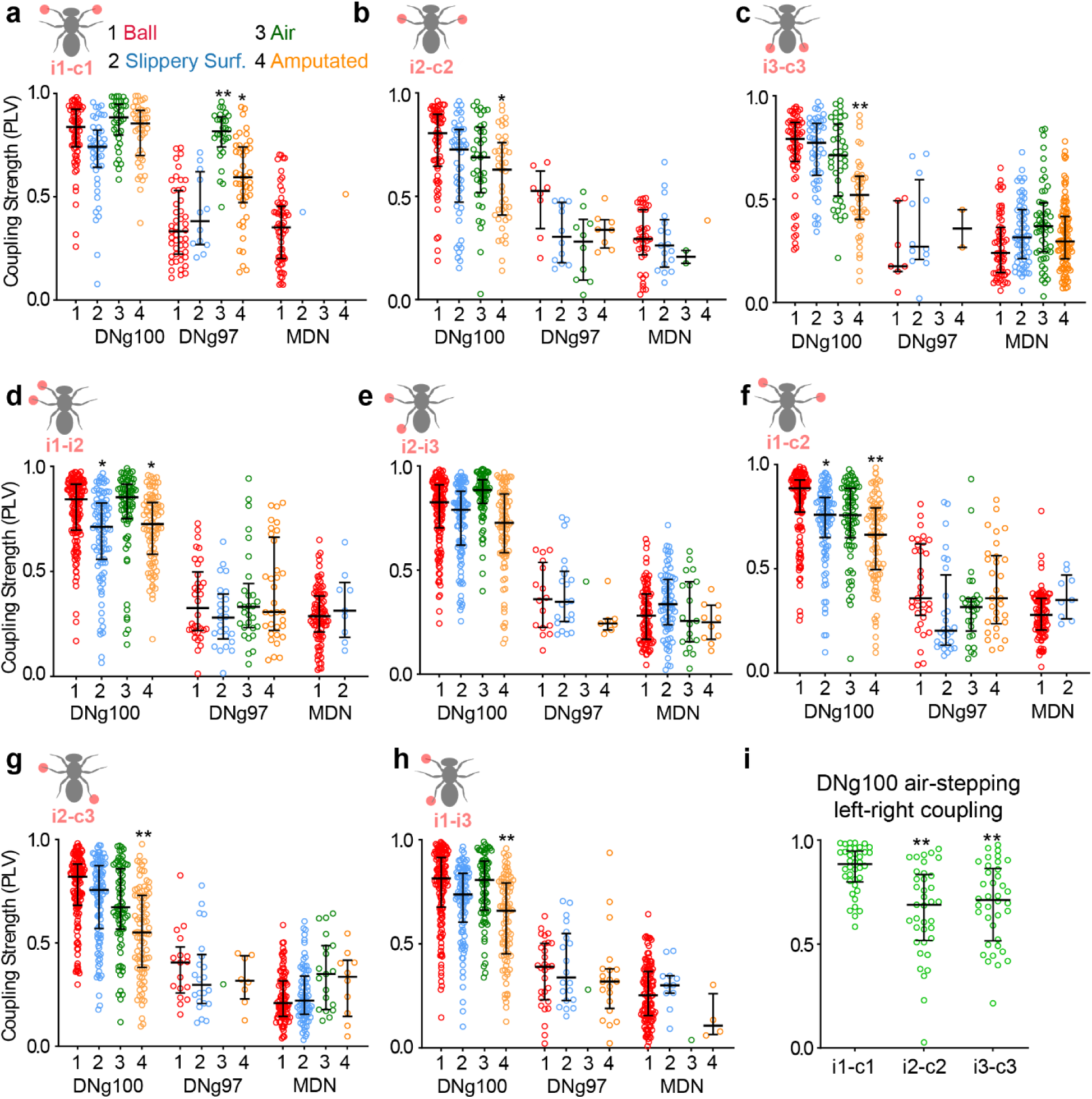
Coupling strength analysis of inter-leg coordination. **a-h.** Coupling strengths corresponding to inter-leg coordination circular phase plots from Fig. 3d-f. n= 28 to 120 trials across 3-12 flies per genotype. Kruskal-Wallis test followed by Dunn’s multiple comparison against condition-1 data within same genotype and condition. (**p <0.001, * p< 0.05). **i.** Coupling strength (PLV) for DNg100 left-right leg pairs in the air condition from panels a-c. Kruskal–Wallis test followed by Dunn’s multiple-comparisons test was performed, with i1–c1 as the reference group (**p <0.001).

**Extended Data Figure 5:**
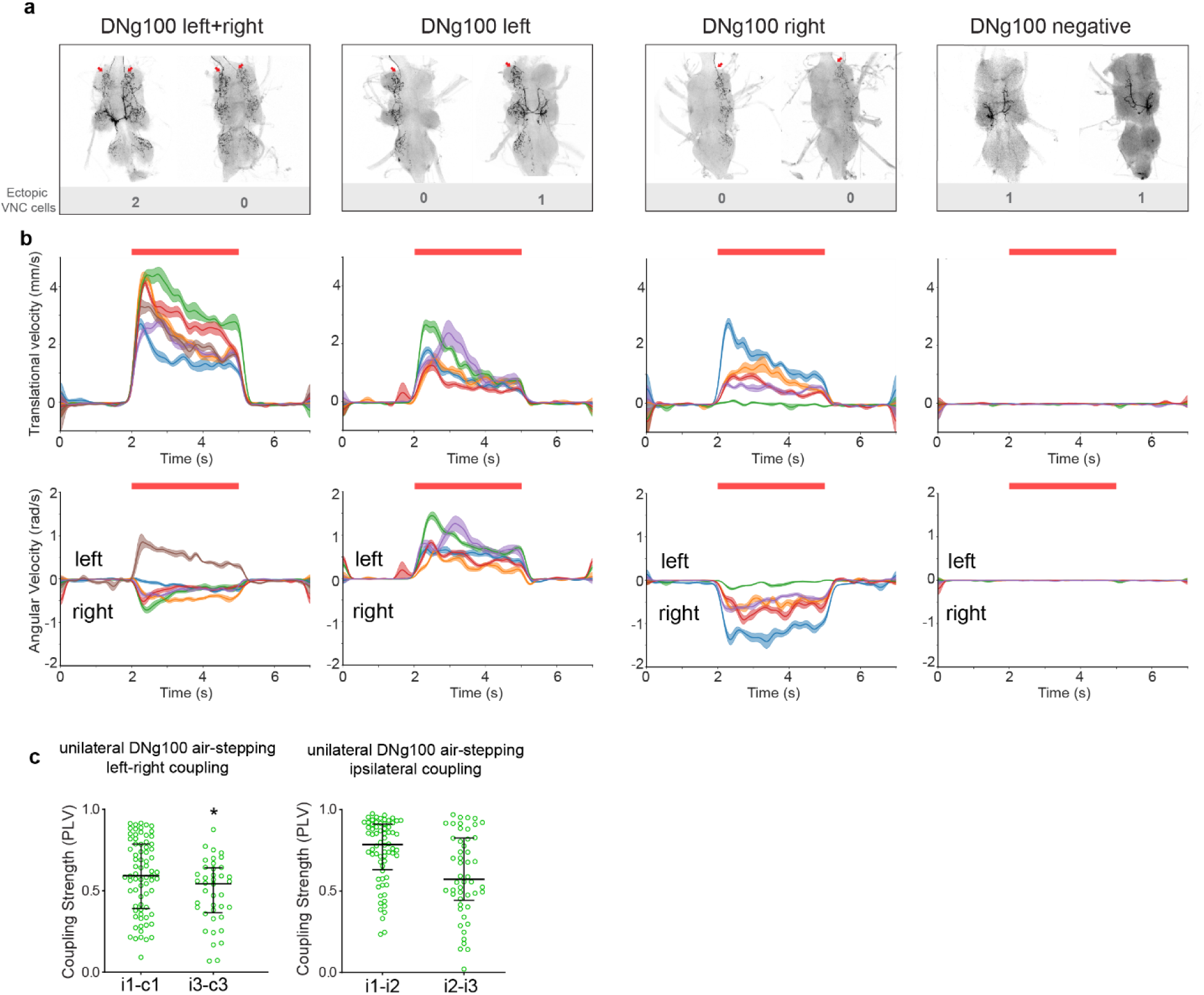
Stochastic activation of DNg100. **a.** Representative IHC images of bilateral (left), unilateral (middle two) and negative (right) classes for DNg100>stop>CsChrimson stochastic labeling experiment. Red arrows indicate labeled DNg100 (unilateral or bilateral) and the numbers in grey indicate the number of non-DNg100 cell types that are labelled in each case. **b.** Forward (top) and angular (bottom) velocity for every fly belonging to each case represented in panel a. The order is the same as panel a. Each trace represents averaged trial velocity per single fly (mean ± s.e.m); red bar on top shows optogenetic stimulation. **c.** Coupling strength (PLV) for unilateral DNg100 left-right leg pairs (left panel) and ipsilateral leg pairs (right panel) in the air condition. N = 92 trials over 10 flies. Mann-Whitney test was performed, with i1–c1 as the reference group for left panel (*p <0.05).

**Extended Data Figure 6:**
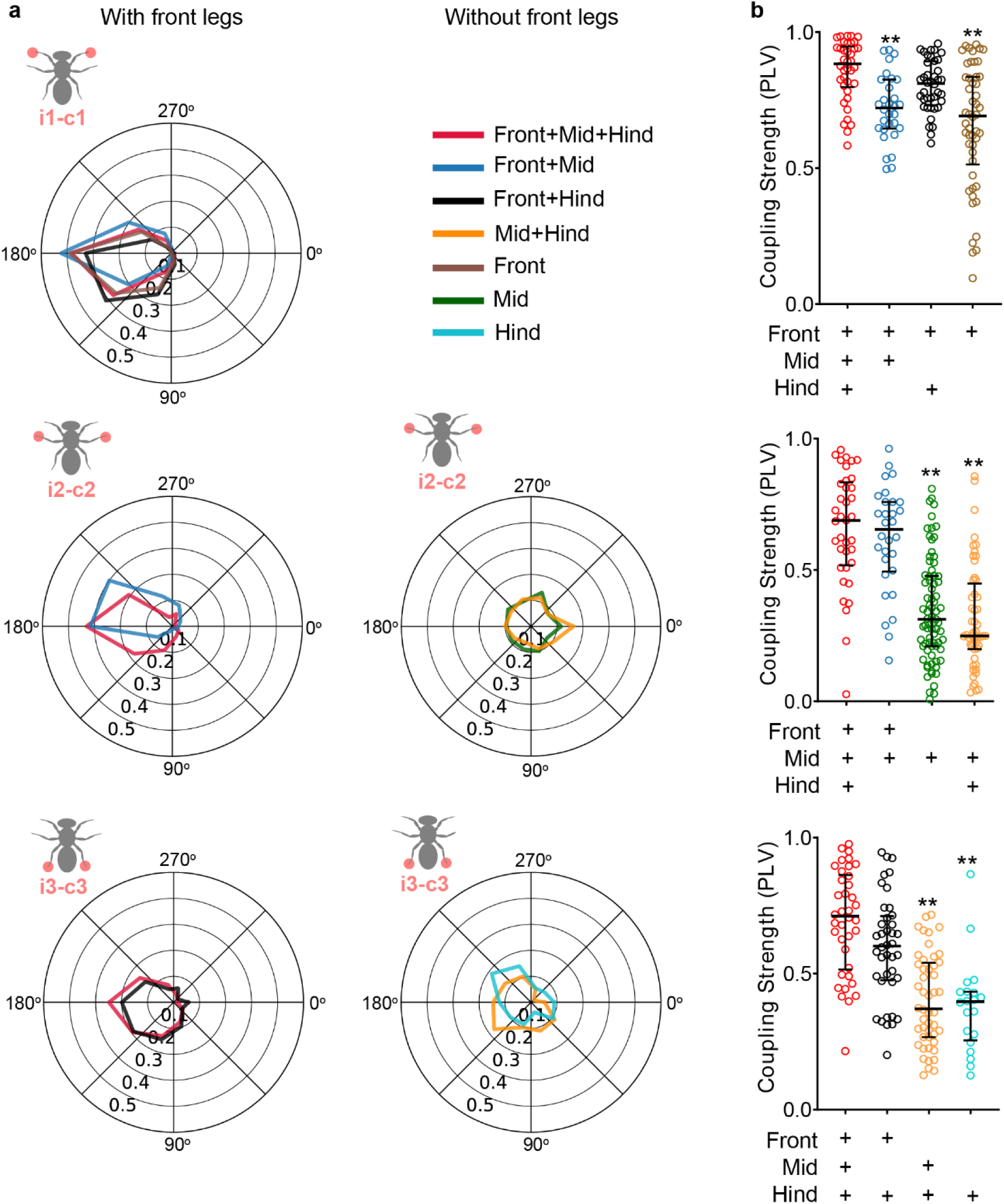
Leg removal experiments in DNg100 stimulated flies. **a.** Inter-leg coordination plots for within-segment contralateral coupling in DNg100 stimulated flies with different sets of legs removed (legend : top right). The front+mid+hind dataset is same as what used in Fig. 3d. Circular phase-plots are organized such that data from flies with front legs intact is in the left column and data from flies without front legs is in the right column, depicting the major impact of front leg removal on mid-leg coupling (i2-c2) and to a lesser extent on hind leg coupling (i3-c3). 20 -78 trials across 2-8 flies. **b.** Coupling strength corresponding to circular plots in the same row. Kruskal-Wallis test followed by Dunn’s multiple comparison using Front+Mid+Hind as control. (**p <0.001).

**Extended Data Figure 7:**
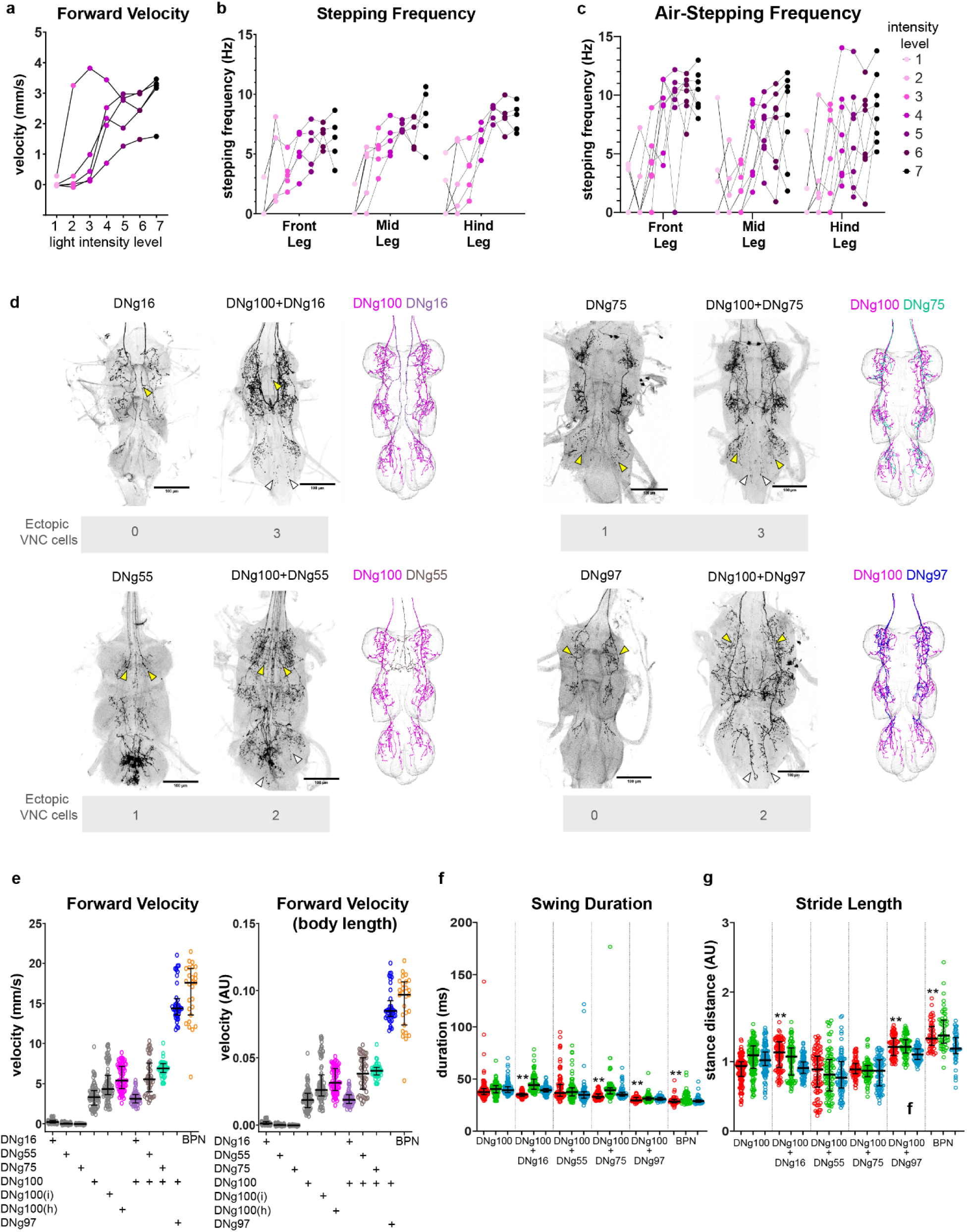
Role of DNg100 and other DNs in speed modulation. **a-c.** Forward velocity (**a**) and stepping frequencies of DNg100 stimulated flies using different light intensities (lowest is level 1 and highest is level 7) in the case of decapitated flies walking on a ball (**b**) or in “air” (**c**). Each curve corresponds to a single fly. **d.** IHC VNC data for DN coactivation in Fig. 4. In each case, empty arrowheads show DNg100 and yellow arrowheads show the additional targeted DN, grey numbers in the bottom indicate ectopic cell types in each case. **e.** Forward velocity in mm/s (left) or normalized to a body length proportional metric (right) for flies with respective neurons activated (reproducing part of Fig. 4b data for comparison and showing normalized velocity). DNg100i (intact flies), DNg100h (homozygous flies i.e. two copies of UAS-CsChrimson; DNg100-Gal4 construct). All flies except for DNg100i and BPN stimulation are decapitated. Stance duration (**f**) and stride length (**g**) corresponding to data in Fig. 4e. Kruskal Wallis test with Dunn’s multiple comparison test with respect to DNg100 stimulation data. Statistical markers only shown for front legs for simplicity. (** p<0.001, * p<0.05).

**Extended Data Figure 8:**
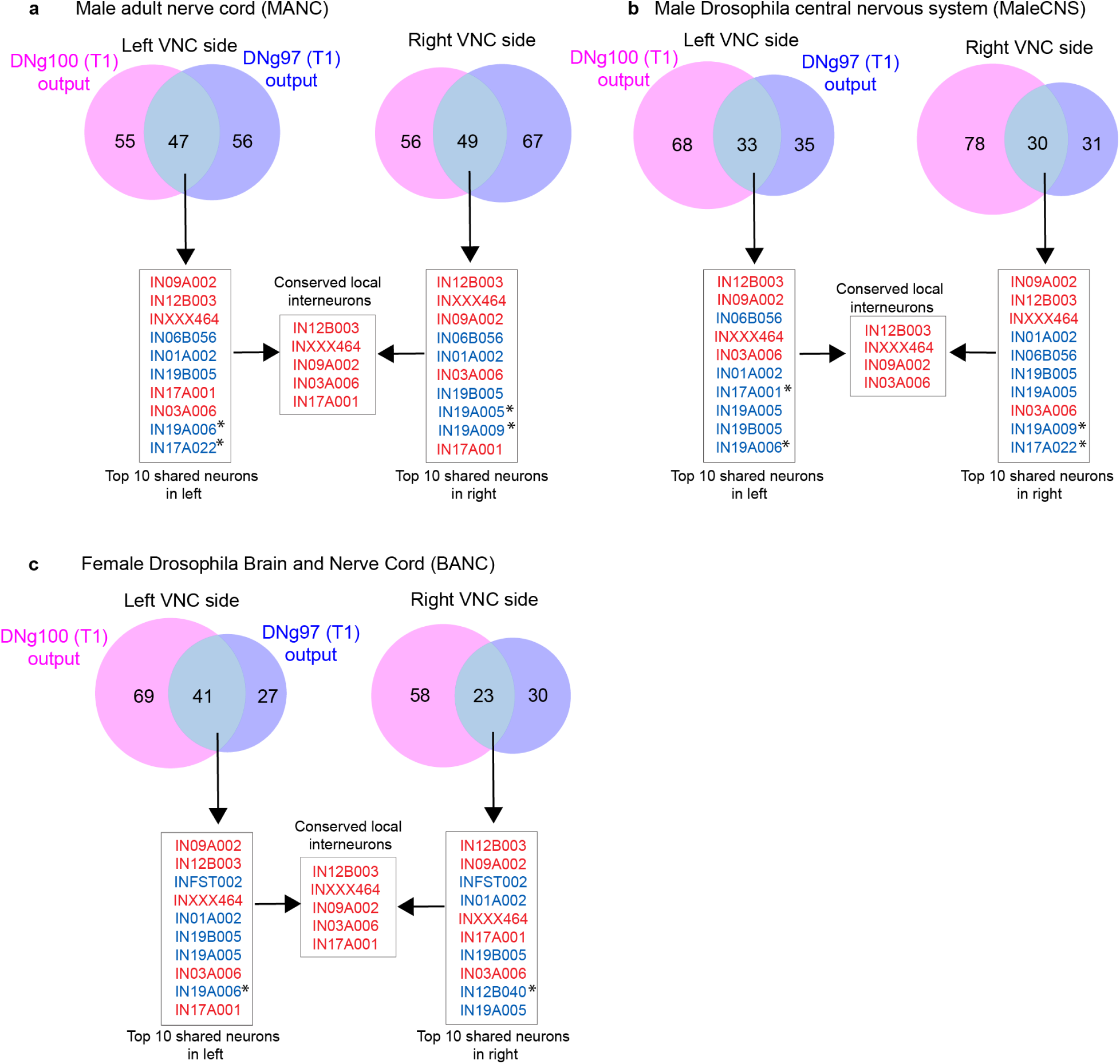
Workflow for selecting conserved CPG motifs across connectomes. **a-c.** Venn diagrams and a flow chart illustrating the selection of CPG motifs from three connectome datasets: MANC (**a**), MaleCNS (**b**), and BANC (**c**). VNC outputs of DNg100 and DNg97 were identified, and shared leg-local interneurons between these two DNs were determined across both left and right sides. In this version, the top 10 shared neurons for DNg100 and DNg97 from each side were selected (shown in red) to define the CPG motif. Asterisks indicate unshared neurons, whereas red neurons denote the conserved shared population identified on both left and right sides and subsequently combined in the merged panel. Note IN17A001 was present in 5 of the 6 hemi-connectomes (only absent in top 10 shared neurons for maleCNS right side potentially due to incomplete synapse numbers) and hence was included in the CPG-motif. Supplementary Table 3 shows all relevant neuron IDs from the corresponding connectome datasets.

**Extended Data Figure 9:**
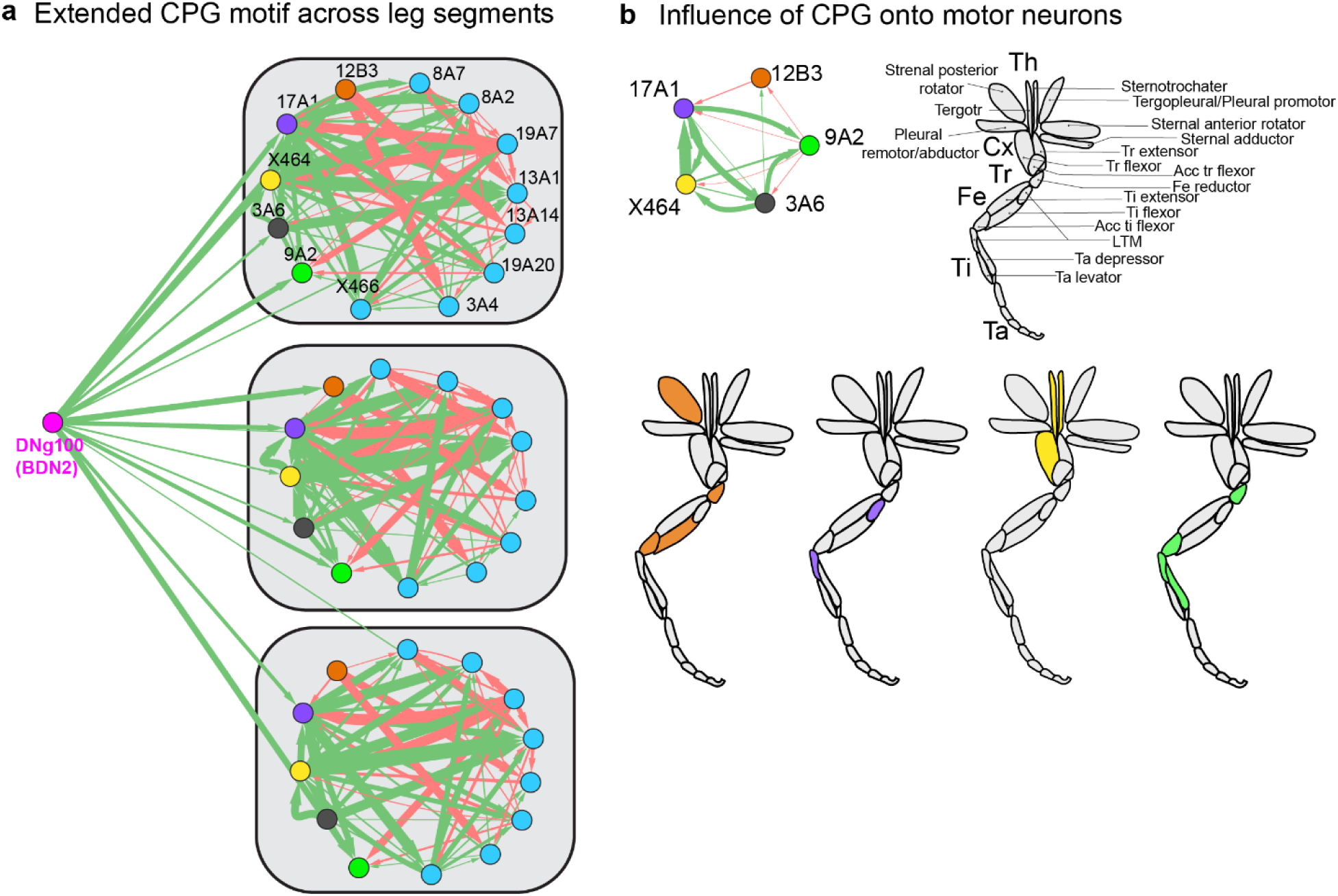
Descending input to the expanded front-leg CPG and CPG influence on motor output from the MANC connectome. **a.** Network from DNg100 to the expanded CPG candidates. Blue nodes indicate neurons added to the core CPG motif to generate the expanded CPG network. (see Supplementary Table 3 for details). Green arrows indicate excitatory connections and red arrows indicate inhibitory connections; arrow width corresponds to synaptic count and scales from 20 to 800 synapses. **b.** Combined unsigned direct and indirect (1-hop) influence of individual CPG neurons on front-leg muscles. Top panel shows the T1 CPG unit and a schematic of the front leg with all muscles labeled. Lower panel shows, front legs with muscles highlighted according to the predicted influence of the corresponding CPG node. For IN03A006, no single dominant muscle target (influence score above an arbitrary threshold) was identified based on influence score. See Methods for influence score calculation. Also see supplementary table 3. Leg schematic adapted from Marin et al.^72^

**Extended Data Figure 10:**
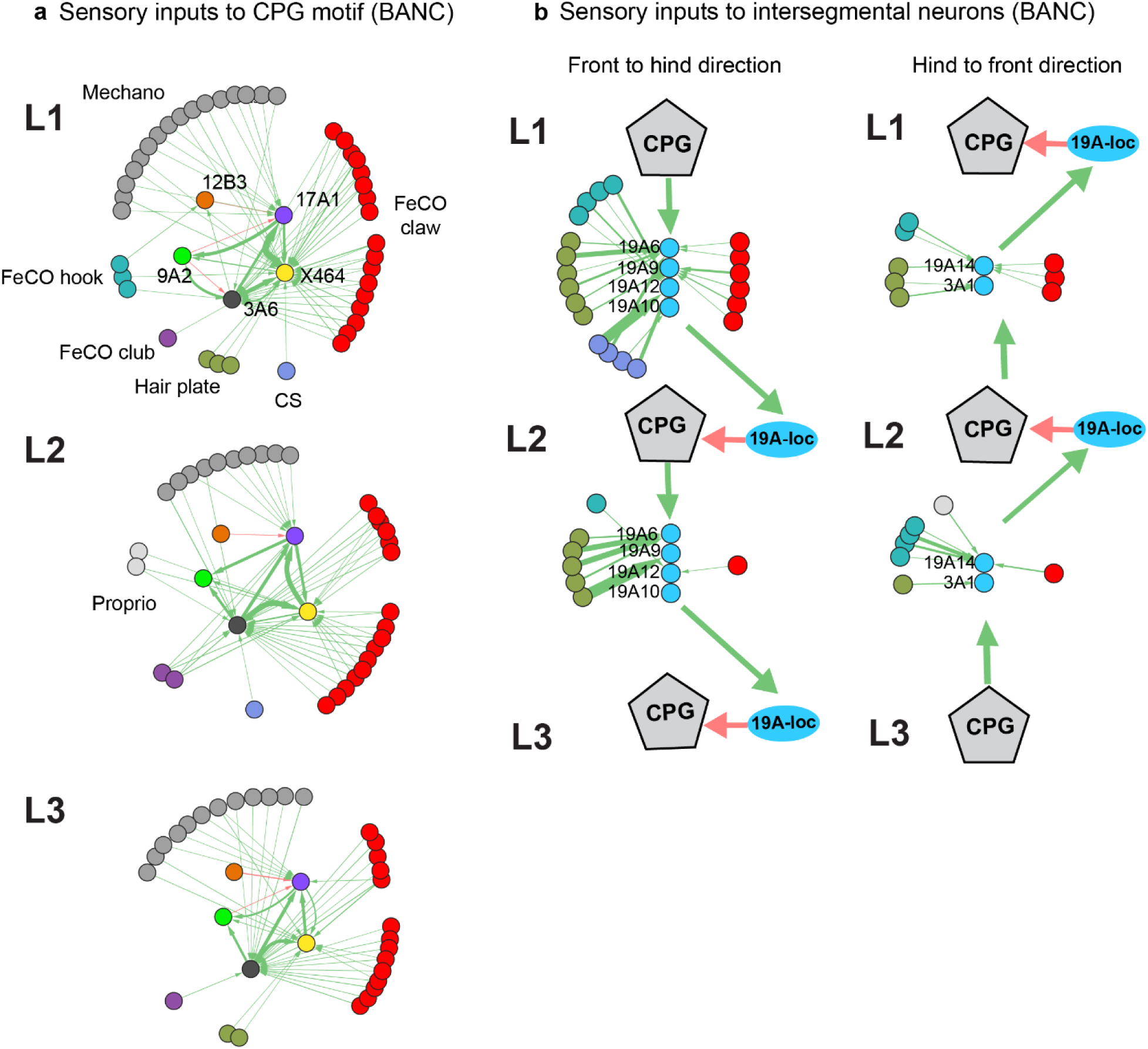
Sensory inputs to CPG motifs and intersegmental neurons in the BANC connectome. **a.** Sensory inputs to CPG motif in front (top), mid (middle) and hind (bottom) leg neuromeres using BANC connectome data which has highest number of annotated leg sensory neurons (among the three available connectomes). **b.** Proprioceptive inputs (hair-plates, FeCO and CS, color-coded as in a) to 19A intersegmental neurons (from Fig. 5f and panel Extended Fig. 10d) identified using the BANC dataset. Green arrows indicate predicted excitatory connections, and red arrows indicate predicted inhibitory connections. Arrow width is proportional to synapse number (scaled from 3-302 synapses in a, and 3-40 synapses in b). Note that the BANC dataset (version v626) have overall higher proofread sensory afferents compared to MANC (version v1.2.3) and MaleCNS (Version v0.9) datasets and hence was specifically chosen for tracing proprioceptive inputs to CPG and inter-leg coordination circuits.

**Extended Data Figure 11:**
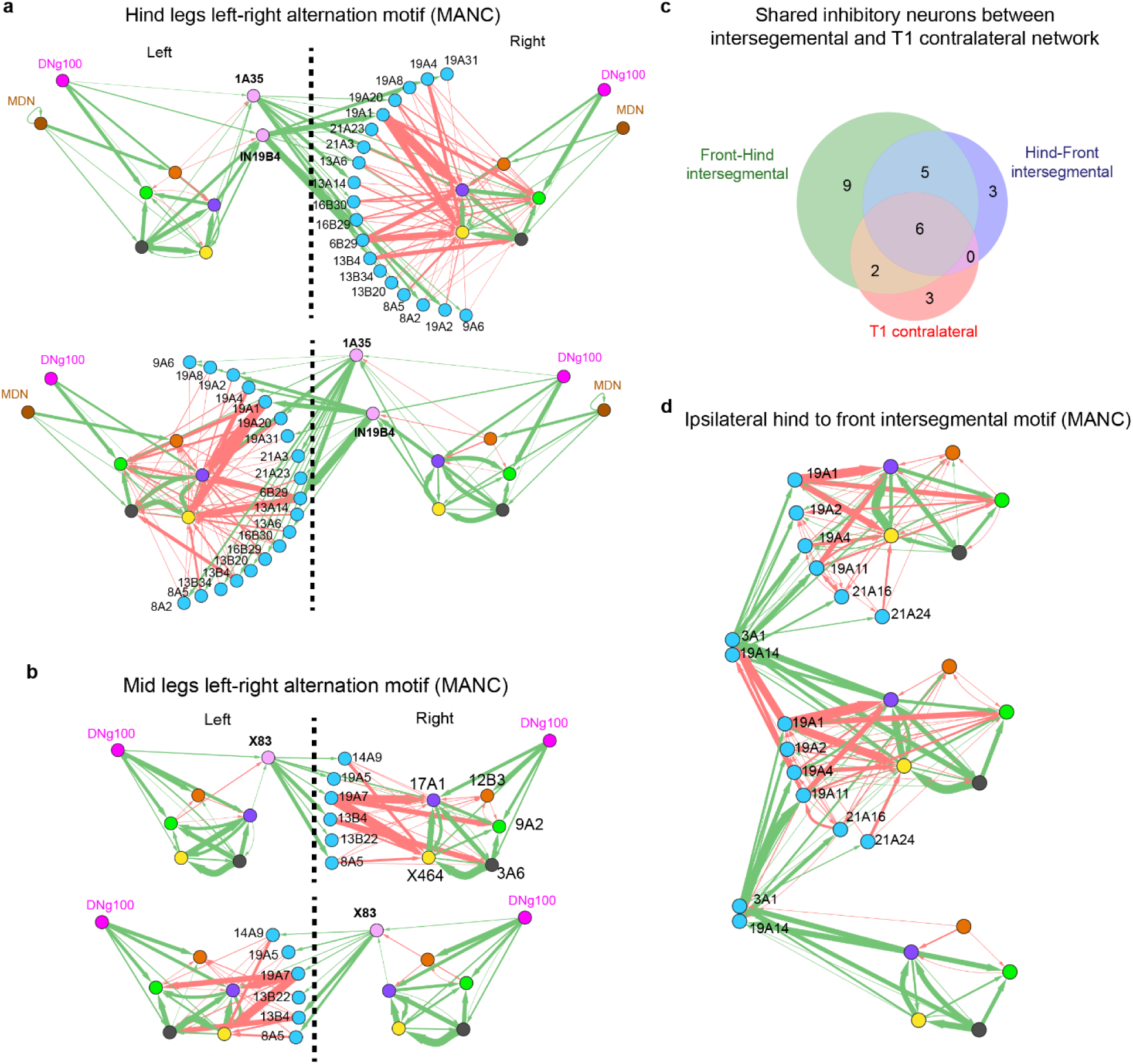
Inter- and intra-segmental connectivity underlying contralateral and ipsilateral coupling. **a-b**. Within-segment contralateral connectivity diagrams for mid legs (**a**) and hind legs (**b**) from the MANC connectome. The dotted line indicates the midline. Neurons shown in light pink represent commissural projections to the opposite side. The top panel in each shows left-to-right connectivity, and the bottom panel shows right-to-left connectivity. **c**. Venn diagram showing the extent of shared local inhibitory neurons identified for within-segment contralateral coupling (using front leg data) and ipsilateral coupling in the front-to-hind and hind-to-front directions. **d.** Ipsilateral hind-to-front network identified using the same approach as the front-to-hind network shown in Fig. 5f. **e**. Green arrows denote predicted excitatory connections and red arrows denote predicted inhibitory connections. Arrow width is proportional to synapse number (scaled from 5 to 637 in a (top), from 5 to 660 in a(bottom), from 5 to 532 b(top), from 5 to 625 in b(bottom) and from 5-946 in d). Also see supplementary table 3.

**Extended Data Figure 12:**
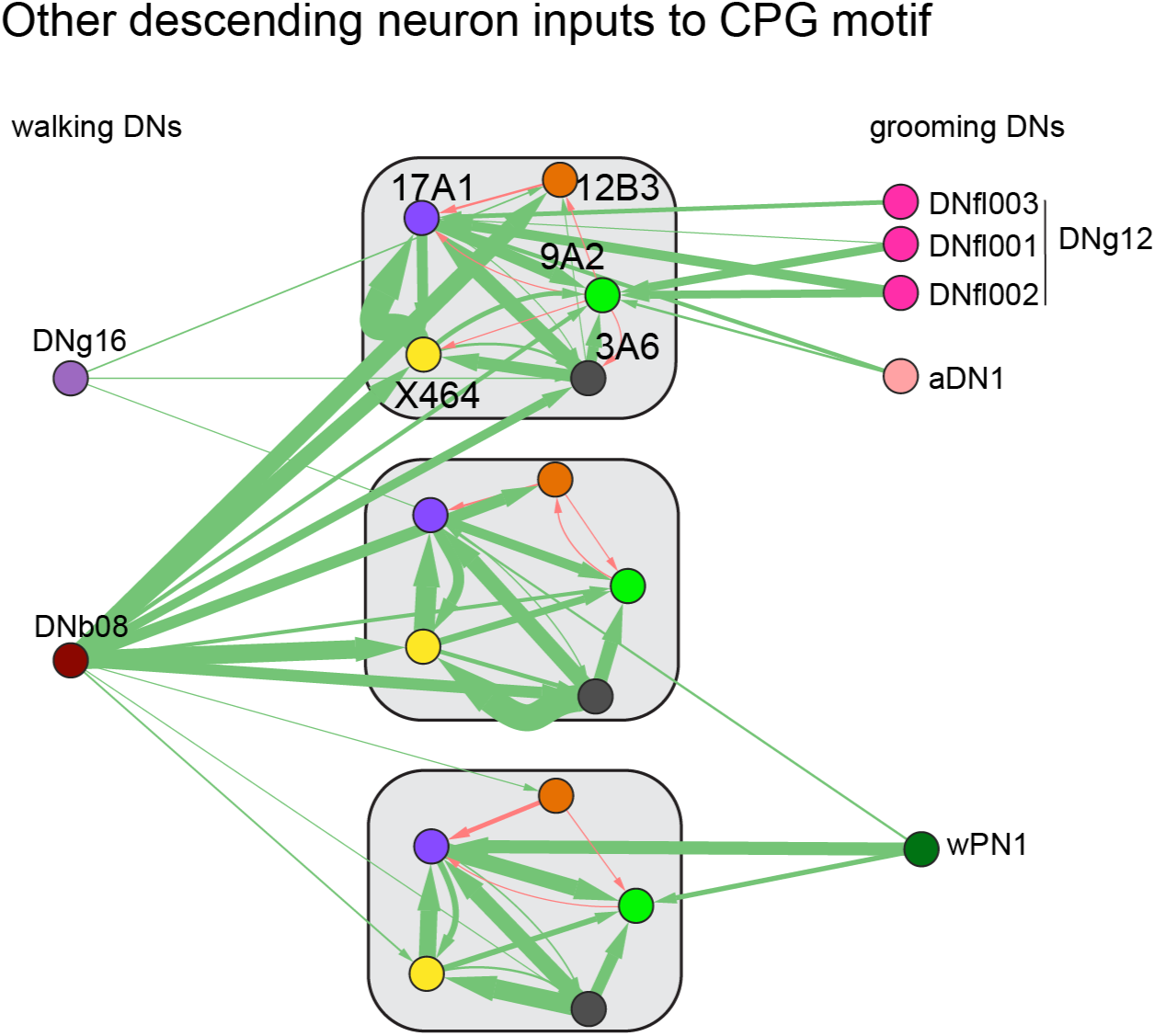
Additional descending neuron inputs to candidate CPG circuits. Connectivity of other descending neurons to the identified CPG network. Walking-related descending neurons not shown previously, including DNg16 and DNb08, are shown on the left. Grooming-related descending neurons, including DNg12 (head and front leg grooming) and aDN1 (antennal grooming by front legs), together with wPN1 (wing-grooming by hind legs), shown on right. Arrow width is proportional to synaptic count (range 5-481). Also see supplementary table 3.

